# Conjugates of neuroprotective chaperone L-PGDS provide MRI contrast for detection of amyloid β-rich regions in live Alzheimer’s Disease mouse model brain

**DOI:** 10.1101/2020.03.08.982363

**Authors:** Bhargy Sharma, Joanes Grandjean, Margaret Phillips, Ambrish Kumar, Francesca Mandino, Ling Yun Yeow, Vikas Nandwana, Vinayak P. Dravid, Xing Bengang, Sierin Lim, Konstantin Pervushin

## Abstract

Endogenous brain proteins can recognize the toxic oligomers of amyloid-β (Aβ) peptides implicated in Alzheimer’s disease (AD) and interact with them to prevent their aggregation. Lipocalin-type Prostaglandin D Synthase (L-PGDS) is a major Aβ-chaperone protein in the human cerebrospinal fluid. Here we demonstrate that L-PGDS detects amyloids in diseased mouse brain. Conjugation of L-PGDS with magnetic nanoparticles enhanced the contrast for magnetic resonance imaging. We conjugated the L-PGDS protein with ferritin nanocages to detect amyloids in the AD mouse model brain. We show here that the conjugates administered through intraventricular injections co-localize with amyloids in the mouse brain. These conjugates can target the brain regions through non-invasive intranasal administration, as shown in healthy mice. These conjugates can inhibit the aggregation of amyloids *in vitro* and show potential neuroprotective function by breaking down the mature amyloid fibrils.

## Introduction

The worldwide social and economic impact of Alzheimer’s disease (AD) dictates the need for the timely invention of suitable solutions for this disease. Failure of anti-AD clinical drugs targeted to neurotransmitters and their association to the toxicity of oligomeric amyloid-β (Aβ) peptides have brought the spotlight back to amyloidosis in AD [1]. The strategy for AD diagnostics needs to be steered towards its early detection using biocompatible diagnostic agents. Magnetic nanoparticles coated with anti-Aβ antibodies can detect pathological amyloid oligomers early in AD brain before they turn into the plaques [2]. Viola *et al.* detected the fluorescently tagged anti-Aβ antibodies in the entorhinal cortex and hippocampus in AD mice brain [2]. They showed that these anti-Aβ antibodies conjugated to magnetic nanoparticles could detect amyloids in AD brain.

Lipocalin-type prostaglandin D2 Synthase (L-PGDS) is the second most abundant protein in the cerebrospinal fluid (CSF) and a chaperone for Aβ peptides [3] and may disaggregate pre-formed amyloids [4]. The composition of the CSF in the brain is very dynamic with a constant turnover of proteins throughout the extracellular and interstitial volumes with 3-5 times daily renewal of CSF at the rate of 0.3-0.4 ml/min in human adults [5]. The protein and metabolite composition of CSF may directly affect amyloid formation in AD brain and reduce the aggregation of amyloids [6, 7]. L-PGDS is intrathecally synthesized in the arachnoid membrane and choroid plexus before getting secreted from astrocytes into the CSF [5, 8]. Its CSF concentration lies within the range of 15-30 mg/l and undergoes circadian changes with serum level fluctuations [5].

In this study to focus on a few aspects of L-PGDS function in AD brain, i.e. whether this abundant neuroprotective chaperone is indeed dynamically associated with the amyloid deposition, what is its turnover or clearance rate from amyloids, and finally explore the feasibility of L-PGDS protein-based magnetic conjugates for magnetic resonance imaging (MRI)-based mapping of Aβ depositions in AD brain. The triple transgenic humanized AD mouse model (3xTg) retains three different transgenes – Presinilin mutation (PS1_M146V_), Amyloid Precursor Protein mutation (APP_KM670/671NL_) (Swedish double mutation, or APP_Swe_), and microtubule-associated tau protein mutation (tau_P301L_) [9]. The deposition of Aβ precedes the formation of neurofibrillary tangles in this model. It was created by microinjecting the embryos harvested from PS1_M146V_ knock-in mice with APP_Swe_ and tau_P301L_ under the control of mouse Thy1.2 regulatory element. Knowledge of L-PGDS localization in AD mouse models can help us better understand its potential role in alleviating amyloid pathology. To detect this, we employ MRI as the imaging tool to identify our conjugated L-PGDS in 3xTg mice brain. Here, we used L-PGDS in conjugation with magnetic nanoparticles for detection of Aβ in AD mice. Intraventricular administration of the conjugates ensures direct circulation through the choroid plexus and diffusion through the brain parenchyma and extracellular space. We observed that administered conjugated L-PGDS penetrate and bind to amyloid structures in AD-associated brain regions despite the abundance of endogenous L-PGDS. Supported by the correlation of AD symptoms with downregulation of L-PGDS concentration in the CSF of AD brain, we posit that L-PGDS targets Aβ in AD mouse brain with high turnover suggestive of fast kinetic binding [10–12]. Therefore, we reason that L-PGDS expressed in other brain regions takes part in other biological functions and can play a more active neuroprotective role. It could mean that a significant pool of brain L-PGDS engages in Aβ interactions which allows the administered exogenous L-PGDS to concentrate into AD- specific brain regions.

MRI is a preferred modality for anatomical and functional imaging due to its additional capabilities in the identification of amyloid-related abnormalities such as edema or hemorrhages [13]. The sensitivity of brain MRI is increased by incorporation of magnetic particles for contrast enhancement [2]. Iron oxide provides a much-needed alternative to the conventionally used Gadolinium-enhanced MRI for AD brain due to recurring reports of nephrogenic systemic fibrosis triggered by its accumulation in patients with renal failure [14]. Iron oxide nanoparticles have been approved as an MR contrast agent by the US Food and Drug Administration, being successfully used to visualize vasculature in clinical cases of brain disorders [15, 16]. Magnetic nanoparticles can be modulated for required characteristics such as biocompatibility, multi-functionality, detectability, and formation of chemical interactions which can suitably be utilized to develop useful clinical agents [17]. The magnetic nanoparticles enriched with inorganic iron shorten the time for spin-spin relaxation perpendicular to the applied magnetic field, or the T2 relaxation, in the proton nuclei surrounding the regions where they localize. T2-contrast enhanced MRI at high magnetic field improves the scanning efficiency and helps to identify the AD brain structures with better temporal and spatial resolution. As a result, there is an enhancement in the negative contrast in T2-weighted MRI, which can be detected as pockets of hypointensity in the images.

## Results

In this study, the magnetic properties of the superparamagnetic iron oxide nanoparticles have been combined with the chaperone function of L-PGDS protein to develop a biomolecular probe with potential for translation into clinical studies of AD. These nanoparticles are functionalized through covalent linkage to bind with recombinant L-PGDS which has an affinity to the aggregated amyloids in AD brain. Binding of the L-PGDS to the nanoparticles can indeed enhance their assimilation into the brain regions [18]. Here we have tested four different iron oxide-based MRI contrast agents for comparison- a) Genetically engineered ferritin nanocages from *Archaeoglobus fulgidus* loaded with 4800 iron (Fe) atoms per spherical protein shell (herein abbreviated as AfFtnAA), b) synthetic magnetic Fe_3_O_4_ nanoparticles with nitro dopamine functionalized Polyethylene glycol (PEG) coating (herein, MNS), c) silica-coated magnetic nanoparticles (herein, SiNP), and d) Dopamine coated magnetic nanoparticles (herein, DNP) [19–21]. A general protocol for the preparation of stable iron oxide nanoparticles involving the decomposition of the iron-oleate complex at high temperatures is reported elsewhere [22]. All the contrast agents used in this study comprise a magnetic core which enhances the MR contrast and a biocompatible outer layer made of biopolymers in synthetic nanoparticles and a hollow protein shell in case of ferritin nanocages. The engineered AfFtnAA used in this study were produced from wild type *Archaeoglobus fulgidus* with alanine residues replacing lysine150 and arginine151 [23]. These nanocages were further optimized for better magnetic properties with enhanced relaxivity of ferritin magnetic core (to be published). PEG coating on the MNS helps the nanoparticles stay longer inside the brain regions by avoiding the process of opsonization by macrophages and therefore, to elicit a lower immune response [24]. Silica coating as the outer layer provides increased biological stability to the SiNP and ease of surface modification to generate multiple functionalizations on the nanoparticles [21]. Dopamine coating enhances the water dispersibility of the nanoparticles, therefore making the DNP more biocompatible [25].

Recombinant wildtype human L-PGDS protein with C-terminal 6x-His tag was expressed and purified from the soluble fraction of *E.coli* BL21 Rosetta cells. The eluted protein was further purified using size exclusion chromatography to get a single purified fraction in the presence of 2mM TCEP reducing agent (Figure S1). 100μM of L-PGDS was used to set up the chemical reaction for the preparation of conjugate probes. The free carboxylic groups on the surface of nanoparticles were covalently conjugated to the free amino groups on lysine and arginine on L-PGDS using carbodiimide crosslinking chemistry [26]. The increase in the size of the conjugated protein was detected from dynamic light scattering (DLS) [27]. The diameter of L-PGDS protein molecules increased from approximately 4nm to 20nm for LPGDS-AfFtnAA (LA) conjugates and close to 100 nm for the conjugates of LPGDS-MNS (LM), LPGDS-SMNP (LS) and LPGDS-DNP (LD) upon crosslinking procedure with overnight incubation (Figure S2). Single peaks in DLS spectra showed successful conjugation of 50μM L-PGDS to the nanoparticles, and no unconjugated species was observed. DLS data collected after a few days of incubation at 4°C showed the propensity of aggregation in LM, LS, and LD conjugates.

To determine any possible neurotoxic effects from the conjugated probes, we incubated differentiated SH-SY5Y neuroblastoma cells for 24 h with different concentrations of L-PGDS protein and all four conjugate probes. Cellular metabolic activity causing NAD(P)H-dependent reduction of 3-(4,5-dimethylthiazol-2-yl)-2,5-diphenyl tetrazolium bromide (MTT) into the purple-colored formazan was used as a marker for cell viability. Addition of concentrations up to 20μM of protein showed no significant toxic effects on cells as measured by absorbance intensity at 570nm (Figure 1-A). The absence of any significant toxic impact on cell viability confirmed the biocompatibility of all the designed conjugates. An interesting trend was observed when the conjugated probes were incubated with the SH-SY5Y cells in the presence of pre-formed Aβ fibrils formed from initial Aβ40 monomeric concentration of 75μM (Figure 1-B). The toxic effects of Aβ on the cells were decreased as the concentrations in the cell culture media were gradually increased. These results taken together imply that all four L-PGDS conjugates have a protective effect against Aβ induced neurotoxicity. On its own, native L-PGDS did not show any significant neurotoxicity towards SHSY-5Y cells (Figure S3).

**Figure 1:**
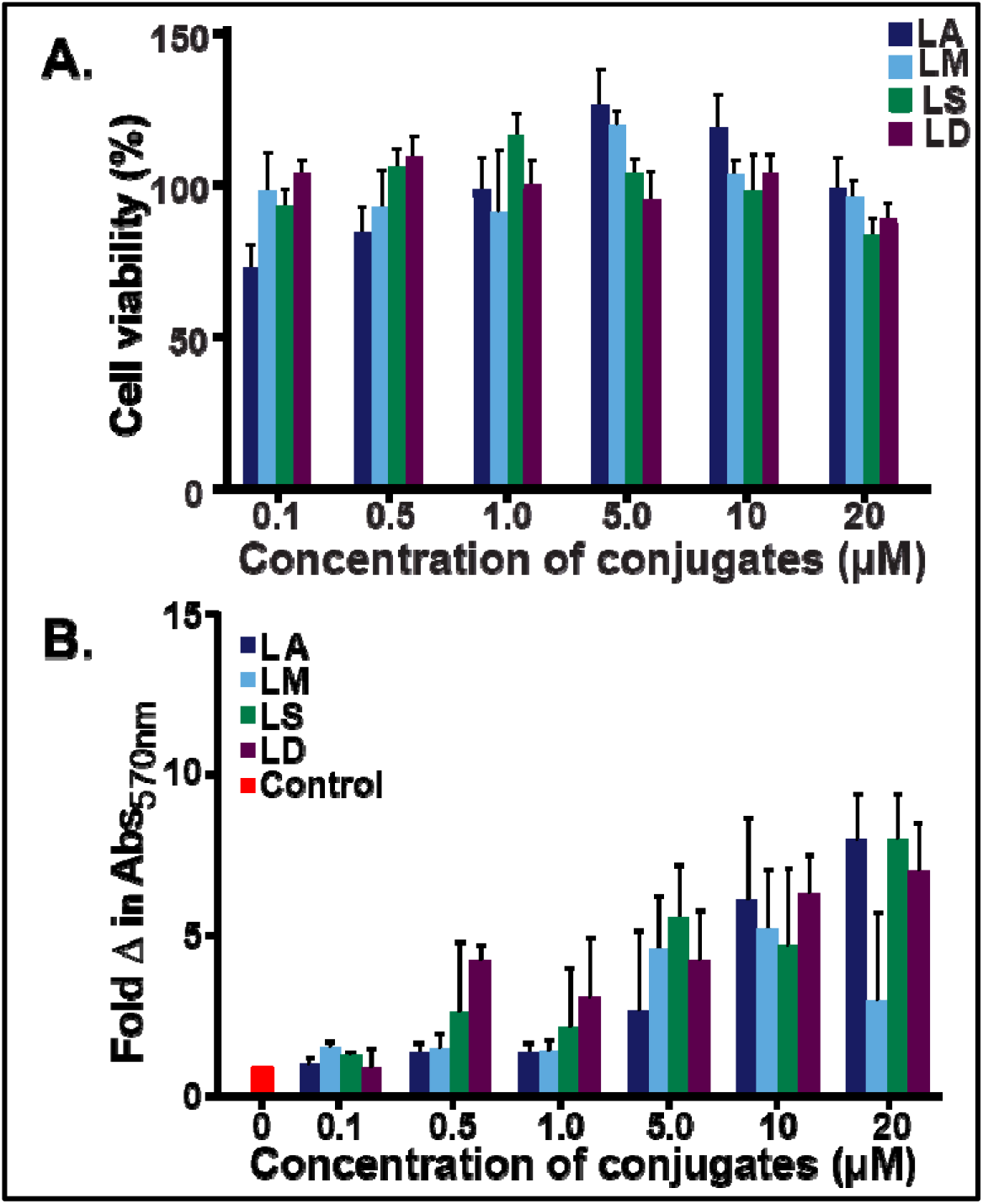
Cell viability of SHSY-5Y cells in the presence of LPGDS-IONP conjugate probes. The mitochondrial reduction of MTT substrate into metabolized formazan product was used to calculate the percentage cell viability of SHSY-5Y human neuroblastoma cells, mean ± SEM (n=3). A. Incubation with different concentrations of the LPGDS-IONP conjugates (LA-blue, LM-cyan, LS-green, and LD-purple) for 24 h. The number of viable cells was correlated to the Abs_570nm_. B. Protective role of LPGDS-IONPs on SHSY-5Y cell growth in the presence of Aβ fibrils.

These four nanoparticles were covalently conjugated with human wildtype recombinant L-PGDS and tested for stability, functionality, and biocompatibility. Contrast enhancement by the four magnetic nanoparticles in T2-weighted MRI at 14T was compared by Rapid imaging with refocused echoes (RARE) based on Carr-Purcell-Meiboom-Gill pulse sequence. The values for *r2* relaxivity were calculated for each nanoparticle and comparing the linear slope. A slope of the intensity versus concentration curve provides a measure of relaxivity (Figure S4). Higher the relaxivity, lower the T2 relaxation time and therefore better the contrast agent. With a hundred-fold increase in the concentration of the nanoparticles, the apparent T2 relaxation time decreased from roughly 150 ms to 50 ms.

The functionality of L-PGDS in the conjugate probes was tested by measuring the enzymatic activity of L-PGDS to generate prostaglandin D2 (PGD2) as an isomerization product from prostaglandin H2 (PGH2). Measurement of the absorbance for the conjugate samples through enzyme-linked immunosorbent assay (ELISA) to quantify the PGD2-methoxime product showed retention of 70% activity in case of protein conjugate LA. The conjugates of L-PGDS with synthetic nanoparticles, LM, LS, and LD showed around 30% activity compared to corresponding concentrations of wildtype L-PGDS protein (Figure S5). The amount of the product was calculated by plotting the absorbance on the standard PGD2-MOX four-parameter curve fit as per the manufacturer’s instructions (Figure S6).

Inhibitory effect of L-PGDS on Aβ40 aggregation as a chaperone was tested for the conjugated probes. The aggregation kinetics of monomeric Aβ40 samples incubated at 37°C was monitored for 72 h in the presence of the L-PGDS conjugates by measuring the fluorescent intensity of Aβ interacting fluorophore Thioflavin T at the 440nm excitation wavelength and an emission wavelength of 484 nm. With due consideration, different sub-stoichiometric ratios of LA conjugates to 25 μM Aβ40 were tested to compare their efficacy as an inhibitor of Aβ aggregation. For 25μM as starting concentration of Aβ40 monomers, 2.5μM L-PGDS conjugates were able to inhibit Aβ40 aggregation (Figure S7). All the conjugated probes were also efficient in inhibiting Aβ40 aggregation, though the efficiency was slightly decreased when compared to unconjugated L-PGDS.

Interaction of LA conjugates on the mature Aβ fibrils was visualized with the help of transmission electron microscopy (TEM). The conjugated probes were localized at the tips along the lateral edges of mature fibrils before breaking them down into smaller fractions (Figure 2). The LA conjugates were able to degrade the mature Aβ fibrils as seen through TEM where co-incubation for 30 min with the Aβ40 fibrils caused the breakdown of mature Aβ into smaller species (Figure S8). The LA conjugates were able to cause up to 80% decrease in the fluorescence intensity of Aβ at an as low ratio as 1:50::LA:Aβ40 (Figure S9).

**Figure 2:**
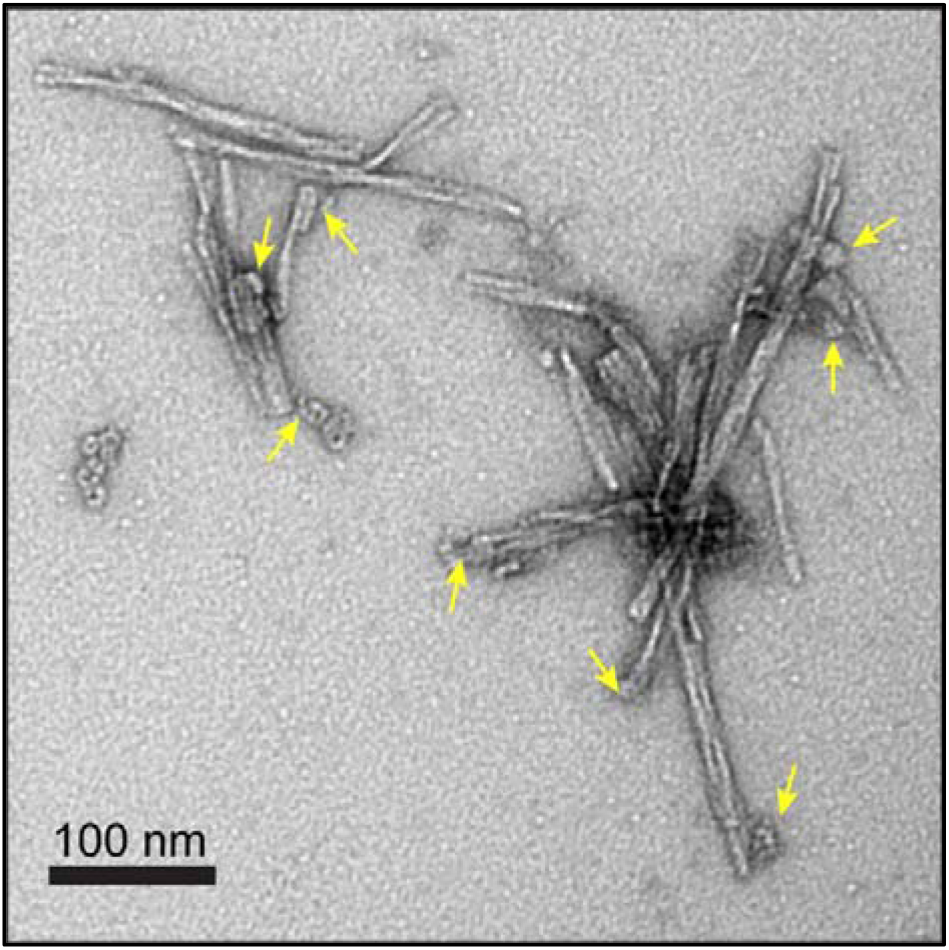
Binding of LA conjugates to the Aβ (1-40) fibrils. Transmission electron microscopy image shows the presence of LA conjugates lateral to the Aβ (1-40) fibrils.

Intraventricular injections of LA conjugates were administered to 1-year old healthy mice and age-matched 3xTg AD mice to see the diffusion of the conjugates towards amyloid-rich regions. 1μl of 50μM LA conjugates were injected into the ventricles of 3xTg AD mice through intracranial injection to discern the distribution pattern through brain regions. LA conjugates administered through intraventricular injections in the 3xTg mice were co-localized with Aβ- rich regions as seen using MRI and histological stains. The localization of conjugates led to hypointensity in these regions as compared to control mice (Figure 3, S10). Compared to the healthy control, hypointensity was observed near the dentate gyrus, corpus callosum, the CA3 hippocampus region and cortical regions such as somatosensory and somatomotor areas, which have been previously shown to correlate with accumulation of amyloids [28–30]. This diffusion through brain regions was confirmed using histological staining of brain amyloids using Congo Red, and conjugated magnetic nanoparticles using Prussian Blue in excised brain slices, which co-localized in the AD-associated brain regions in the brain slices excised after 72 h (Figure 4, S11). LA conjugates injected through intracranial injections into the ventricles in 3xTg AD mice, with similar Aβ plaque formation as humans, were seen largely accumulated into the choroid plexus before diffusion into other brain regions [31]. With the progression of time to 72 h post-administration, the regions of hypointensity and susceptibility variations gradually propagated to amyloid-rich regions including hippocampus and cortical nuclei as seen in gradient recalled echo (GRE) T2* images (Figure 5, S12). Although the GRE sequence is more sensitive to T2* changes in MRI, there is an inherent lower signal-to-noise ratio as compared to spin-echo sequences which often provide high-resolution T2 images (Figure S13). Since the GRE sequences use a low angle shot, the echo amplitude is often too low, which attributes to this limitation [32]. We observed hypointensity in cortical nuclei, hippocampal regions and other AD-associated brain regions with a decrease in T2 observed for 72 h post intraventricular injections (Table S1). AD-associated anatomical changes are known to progress from the entorhinal cortex to other cortical regions via hippocampus [33]. Since the conjugates directly targeted the amyloid rich regions in the AD mice brain, we posit that the presence of recombinant L-PGDS conjugates does not affect the turnover of intrathecal L-PGDS endogenously present in the brain. GRE images show that most of the administered conjugates reached the amyloid-rich brain regions including corpus callosum, thalamus, cortex and cerebral nuclei within 48 h as observed through increase in hypointensity in these regions and did not diffuse out of these regions, except the hippocampus, till 72 h post-injections (Figure 6). Therefore, through intraventricular injections, 1 μM of administered L-PGDS can easily diffuse to major AD-associated brain regions within 48 h.

**Figure 3:**
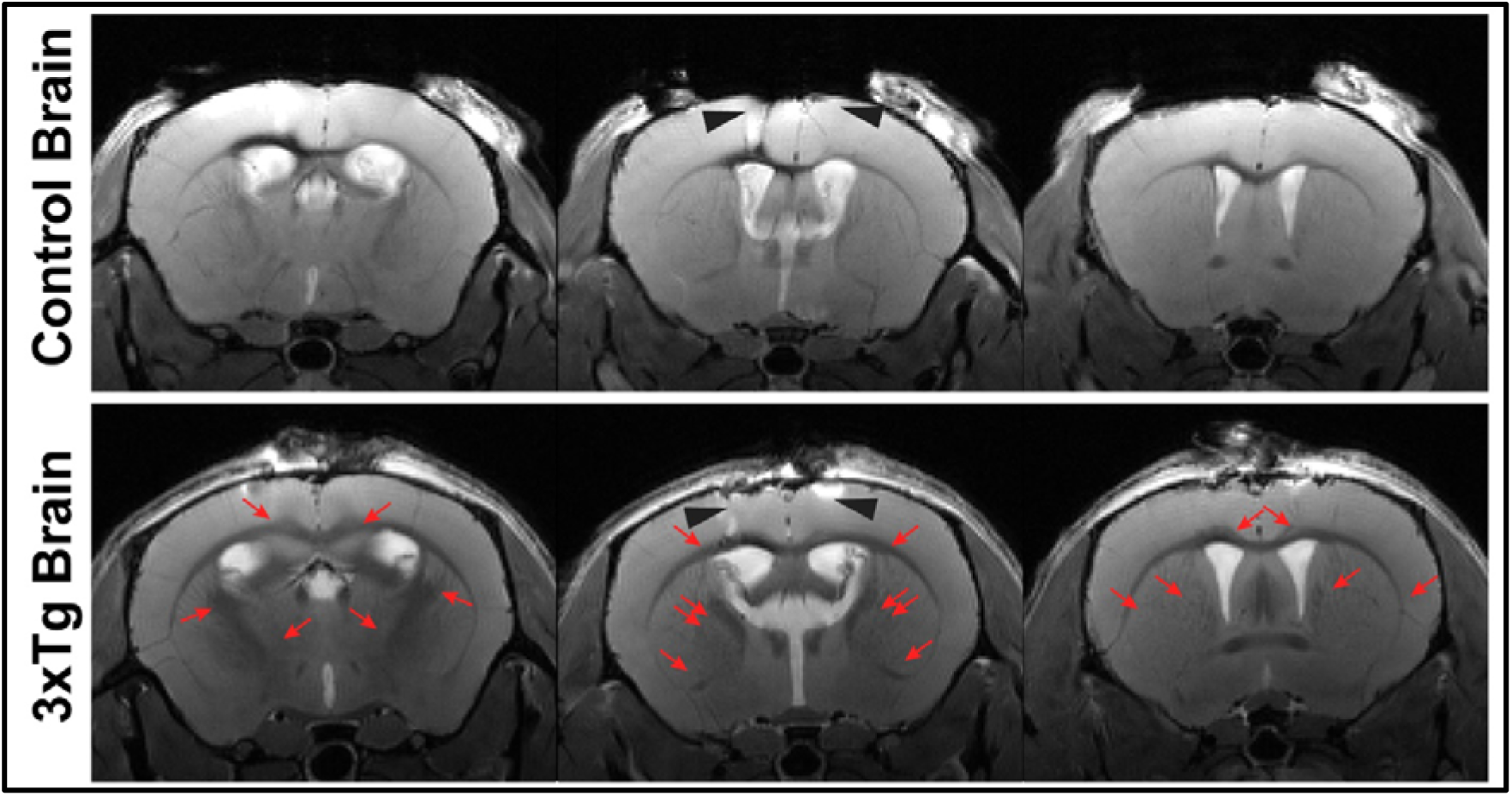
Contrast comparison upon intraventricular injections of LA conjugates in healthy versus AD mice brain. Enhancement of T2* contrast in 1-year old 3xTg mouse brain compared to age-matched control mouse brain. The increase in hypointensity is marked by red arrows in the 3xTg brain. Black arrows in the middle images in both panels mark the entry point of intraventicular injections. The coronal images were acquired at 11.4 T using T2_TurboRARE sequence with image size was 200 × 106 pixels, with field of view 17 × 9 mm, RARE factor: 4, TE 15.56 ms, TR 3500 ms with 38 slices of thickness 0.4 mm each through the mouse brain, scan time: 3 min 2 sec, motion averaging, flip back, fat suppression. We have provided all slice panels without arrows in the supplementary.

**Figure 4:**
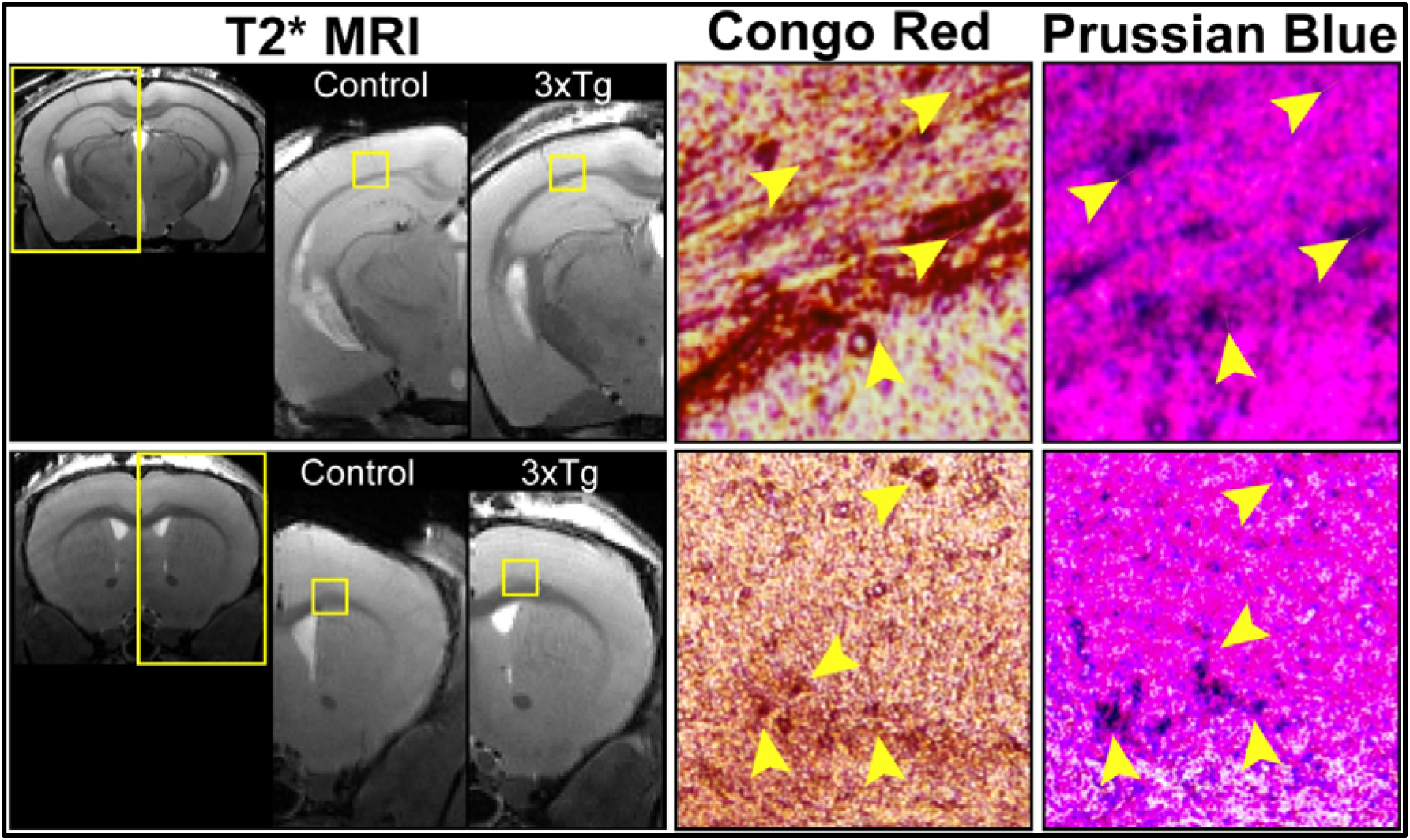
Histological staining of 3xTg AD brain slices post intraventricle injections. Regions of hypointensity in T2* MRI images are stained with Congo red and Prussian blue staining of mouse brain slices of different regions after excision from 3xTg mice 72h post intraventricular injection with LA conjugates. More slice panels are provided in the supplementary.

**Figure 5:**
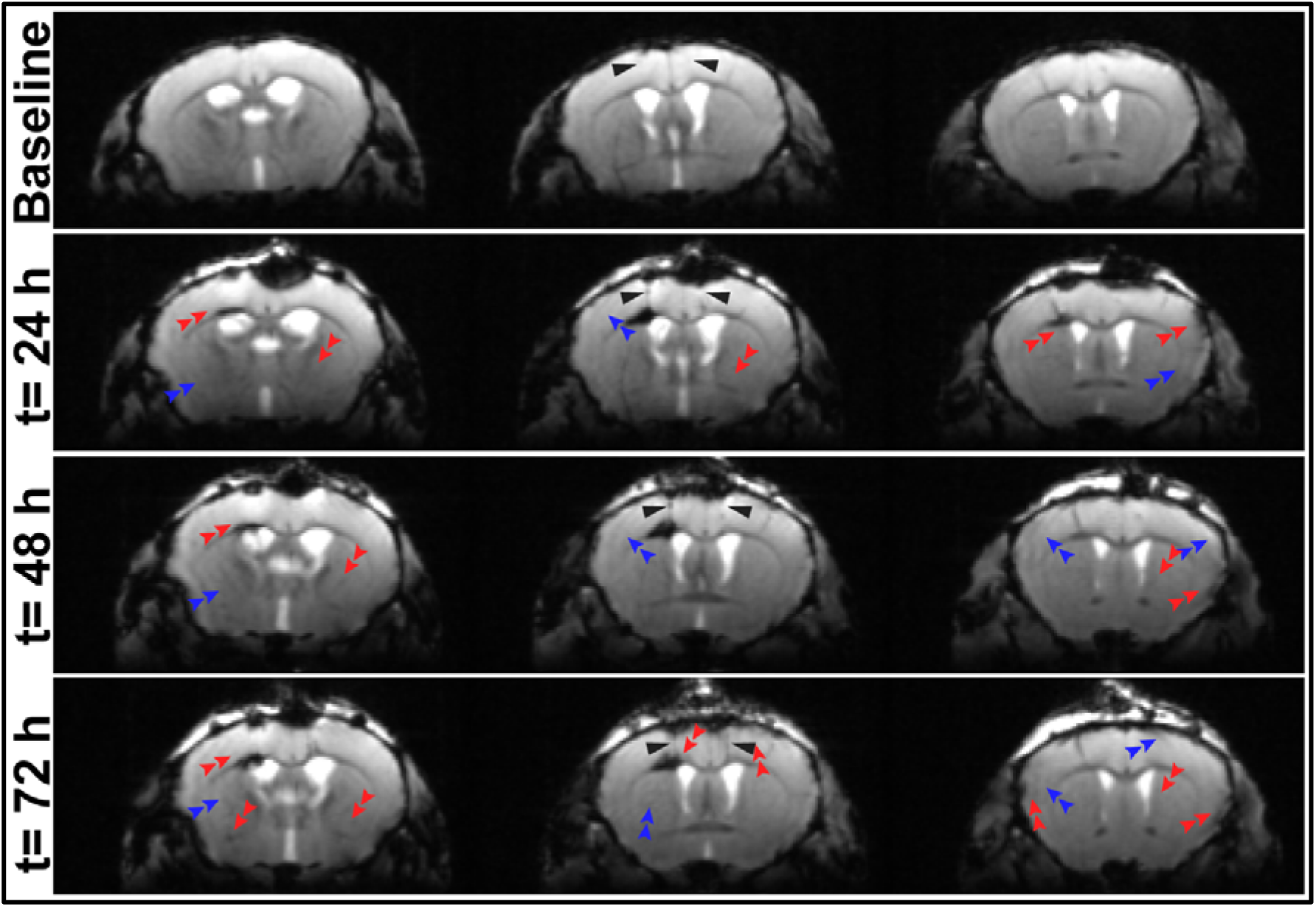
Comparison of T2* contrast enhancement in the 3xTg AD mouse post intraventricular injection of LA conjugates. Regions of hypointensity (red) and susceptibility (blue) variation due to the iron deposits in the mid-brain slices seen in the images 24 h, 48 h, and 72 h post intraventricular injection images have been marked with red arrows. Black arrows in the middle images in all panels mark the entry point of injections. The coronal images were acquired at 11.4 T using T2*-FLASH sequence with an image size 200 × 106, field of view 17 × 9 mm, Flip angle 60°,TR: 1500 ms, TE: 10 ms, for 38 slices with thickness 0.4 mm each through the mouse brain; scan time: 6 min, motion averaging, fat suppression. We have provided all slice panels without arrows in the supplementary.

**Figure 1.**
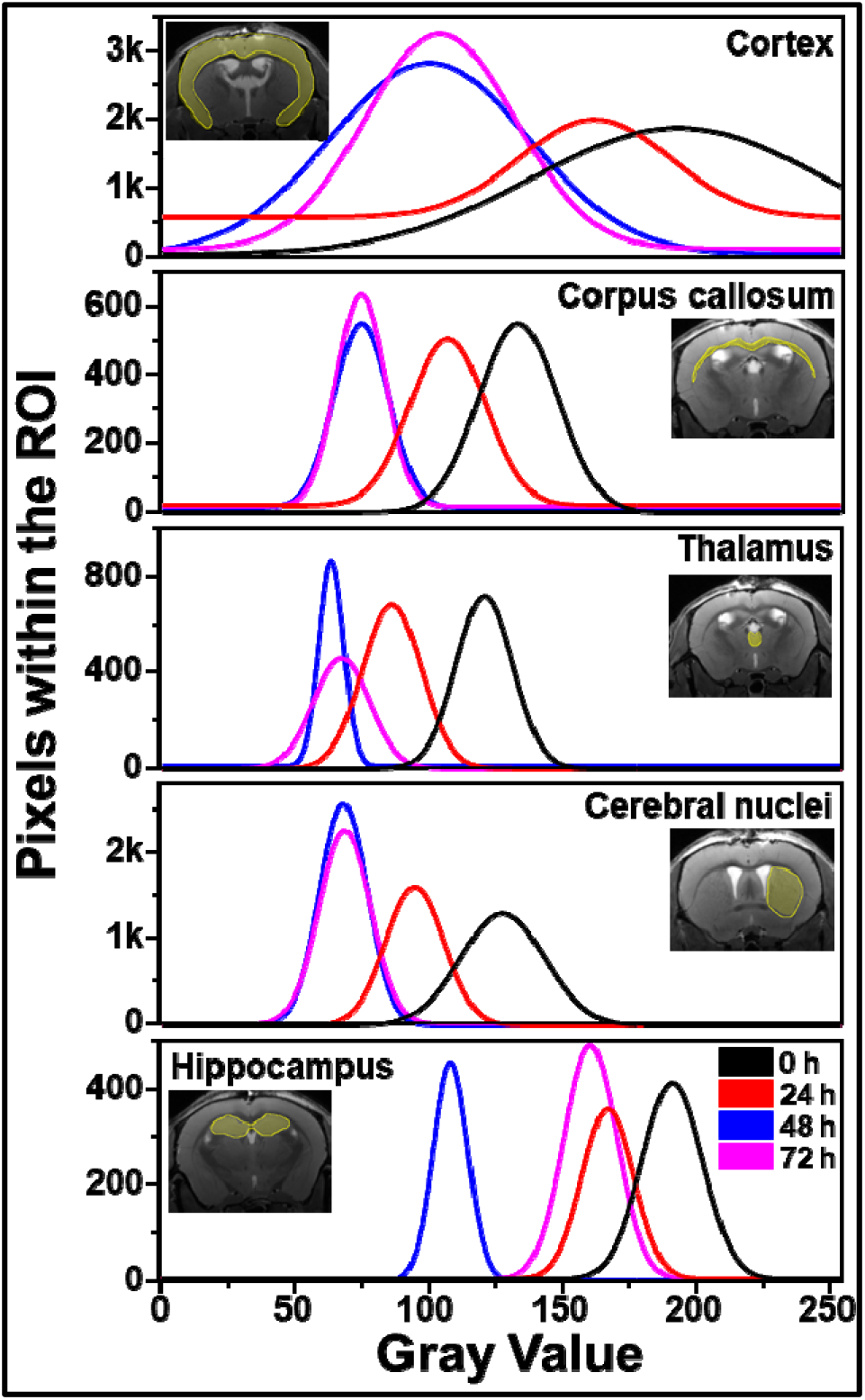
Mean values of grayscale pixel value for T2* MRI images post intraventricular injections into the 3xTg mouse brain. The curves show the fit for mean value in different brain regions of interest (ROI) associated with AD as observed in the T2* weighted images taken over a period of 72 h. The curves are shown for just before injections (0 h, Black), 24 h post injections (red), 48 h post injections (blue), and 72 h post injections (pink). The corresponding ROIs are highlighted in the inset of each graph.

We also tested the possibility of targeting the probes through intranasal administration for their transport to the brain regions via olfactory or trigeminal pathway [34]. Non-invasive intranasal administration of conjugates can facilitate direct entry to the brain by-passing the blood-brain barrier (BBB), which poses as a stringent barrier during systemic transport. Following the pioneer study for intranasal administration of conjugated nanoparticles by Viola *et al.*, we show that intranasal administration of these LA conjugates can ensure direct targeting of brain regions such as the hippocampus [2]. 5 μl of LA conjugates were administered through each nostril to anaesthetized J6 control mice. T2* MRI scans post 4h versus 24 h of administration showed enhanced hypointensity in the brain regions near the olfactory bulbs (Figure 7, S14). These alternative modes of probe administration have been explored to use nasal insulin as a possible treatment for AD targeting the insulin receptors in the brain [35]. These observations were confirmed by histology using the Prussian Blue stain which showed the presence of L-PGDS conjugates in the isocortex and hippocampal regions, in addition to the olfactory lobes (Figure S15). The decrease in T2* relaxation in the healthy mouse brain after intranasal administration indicates the successful transport of the conjugate into the brain. This non-invasive method of brain targeting by conjugated L-PGDS probes can open new avenues in the field of AD diagnostics.

**Figure 7:**
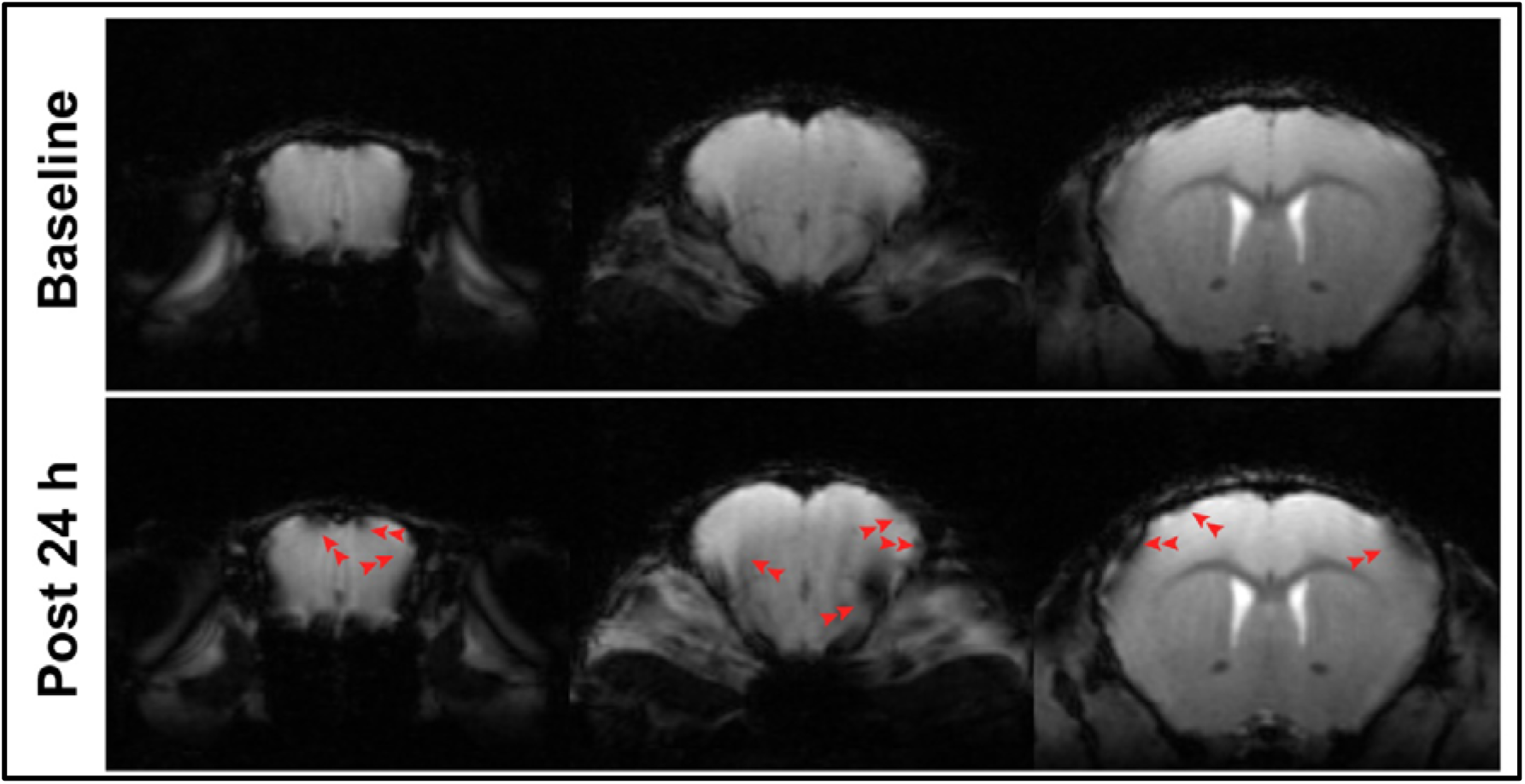
Brain detection of conjugates in control mice by intranasal administration. The left and right panels represent the same slice in T2*-weighted FLASH images taken before and 12 h after the intranasal delivery of LA conjugates in the healthy mice. The coronal images were acquired at 11.4 T using MSME scans with image size was 200 × 106 pixels, with field of view 17 × 9 mm, 60° flip angle, TE 10ms, TR 1500 ms with 3 averages each for 38 slices through the mouse brain. We have provided all slice panels without arrows in the supplementary.

## Discussion

The cost of production of monoclonal antibodies and the requirement of regular administration is a significant drawback of the existing immunotherapies tested for AD treatment [13]. Five of the nine vaccines in clinical trials for AD target different forms of Aβ [1]. The production costs of humanized anti-Aβ antibodies like Solanezumab and Bapineuzumab is very high, especially considering the little significance in their impact on AD patients and a possible list of side-effects as the brain tries to defend itself from a foreign molecule. Being an endogenous brain protein, L-PGDS overcomes the immunoreactivity barrier, and the cost of production is much lower than raising monoclonal antibodies. Our conjugates provide sufficient contrast to develop potent diagnostic tools which can help identify the turnover rate of soluble amyloids into plaques. Intranasal administration of functionalized nanoparticle conjugates expedites their targeting to the AD brain [2]. L-PGDS inhibits the aggregation of Aβ fibrils and acts as an extracellular Aβ disaggregase, not relying on the hydrolysis of Adenosine triphosphate (ATP) for its action [4]. Its key role in the sleep-wake cycle, differential expression in neuronal cells, upregulation in neurological disorders, and interactions with the receptors in the brain make it an ideal candidate for the AD probe design [36–39]. Furthermore, L-PGDS has been previously used to transport small lipophilic drugs into the brain and formulated in brain medications to enhance their targeting across the BBB [18, 40]. We showed that L-PGDS is capable of binding more than 20 proteins from the insoluble aggregates in human AD brain [4]. It forms a stable complex with Aβ40 monomers to inhibit their aggregation and binds to the mature fibrils triggering their disaggregation into smaller species. L-PGDS has been actively used in drug delivery systems due to its chemical behavior as a non-toxic and non-immunogenic transporter of small lipophilic molecules [41]. It mediates neuroprotection by facilitating the clearance of biliverdin by-product in cases of aneurysmal subarachnoid hemorrhage and scavenges harmful reactive oxygen species to prevent neuronal apoptosis [40, 42]. L-PGDS plays a vital role in the sleep-wake cycle, and the inhibition of its activity is associated with sleep deprivation [43]. It is also reported that the Aβ accumulation is triggered by sleep deprivation, and there are implications that alterations in sleep patterns may modulate Aβ pathology [44]. It is diagnostically valuable as a clinical marker for CSF fistula and normal pressure hydrocephalus [45, 46]. Here, we show that L-PGDS can be reliability exploited as a diagnostic tool for AD by its capability as a chaperone and transporter protein accessible to the brain regions. With an easy preparation, this strategy can be used for protein-based drug formulations, which are biologically superior to the antibody-based drugs. The Aβ-chaperone function of L-PGDS was studied by Kanekiyo *et al.*, where they found L-PGDS to co-localize with plaques in murine as well as the human brain [3]. Here, we demonstrated the diagnostic potential of L-PGDS in live AD mice using non-invasive modality like MRI.

This work is a proof-of-concept study to identify biocompatible formulations of protein conjugated with contrast agent which can be used in early diagnostics of Alzheimer’s disease. With this probe based on endogenous brain protein L-PGDS, we have mediated the use of the inherent Aβ chaperone activity of the protein in combination with the detection capabilities of MR active contrast agents. We showed the efficiency of these conjugates as diagnostic agents through MRI studies in anesthetized mice. Additionally, the *in vitro* studies show the therapeutic potential of L-PGDS conjugates in breaking down the mature amyloid fibrils and demonstrate the protective impact on neuronal cells to alleviate the toxic effect of amyloid. Experiments conducted on larger cohorts of mice are required to establish the full potential of L-PGDS as a theranostic tool for early diagnosis of AD. Longitudinal monitoring to study the long-term impact of administrating these conjugates to 3xTg mice in comparison with untreated animals can identify the clinical translational outlook of these probes. Studying a larger population would allow the MRI protocols can be adjusted with proper calibration to contrast differences between age-matched control and AD mice over time. The formulations of the magnetic nanoparticles can be further engineered towards achieving higher *r1* and *r2* relaxivity to render better contrast. Additionally, the issues arising due to magnetic susceptibility artefacts which interfere with the elucidation of the results can be overcome by combining MRI with other safer detection modalities such as ultrasound and SPECT which can, in turn, provide corroboration for the overall impact of administering these molecular probes to the diseased mice. At the same time, other aspects of AD pathogenesis need to be further explored in this quest for the earliest possible detection. Target proteins for other biomarkers of AD such as tau of the characteristic neurofibrillary tangles, chitinase-3-like protein 1 which is linked with inflammation, and the neurofilament light which is indicative of axonal injury are potential avenues which can be explored for AD theranostics [47]. Our results pave the way for the study of other endogenous proteins with significant potential as a chaperone for aggregation-prone amyloids [48]. This work highlights the importance of the use of modifiable contrast agents with engineered coatings to enhance detection of properties of biocompatible diagnostic probes.

## Materials and Methods

### Protein purification

Recombinant human wild type L-PGDS was overexpressed in BL21-DE3 Escherichia coli bacterial cells using previously published protocol [49]. The protein eluted after immobilized metal affinity chromatography (IMAC) using complete His-tag purification resins (Roche) was concentrated using 10KDa cut-off amicon filters (Merck) before further purification. The concentrated protein was passed through Superdex 75 10/300 GL size exclusion chromatography column (GE Lifesciences) to get pure protein fractions in buffer containing 20mM (4-(2-hydroxyethyl)-1-piperazineethanesulfonic acid (HEPES), 150mM sodium chloride (NaCl) and 2mM tris(2-carboxyethyl)phosphine (TCEP) at pH 7.4. Purified protein fractions showed a single band around 20KDa by Sodium Dodecyl Sulphate polyacrylamide gel electrophoresis (SDS-PAGE).

### Conjugation

1-Ethyl-3-(3-dimethylaminopropyl) carbodiimide (EDC) crosslinkers were used to activate the carboxyl groups on the surface of nanoparticles which were stabilized for conjugation using N-hydroxysulfosuccinimide (sulfo-NHS), both stocks freshly prepared in 2-(N-morpholino) ethane sulfonic acid (MES) buffer. All contrast agents were activated for bioconjugation with purified L-PGDS using by addition of 100mM EDC and 250mM sulfo-NHS dissolved in amine- and carboxyl-free MES buffer (Thermo Scientific™) with pH adjusted to 6.0 to a final ratio of 1:5 [50]. Post 1 h of incubation with constant shaking at room temperature, 50μM L-PGDS was added to each CA upon activation. The reaction mixture was incubated at room temperature overnight with gentle shaking. The excess of free carboxyl groups was quenched by adding 1M Tris buffer with pH adjusted to 7.4.

### Dynamic Light Scattering

Dynamic light scattering (DLS) was used for size calculation of L-PGDS conjugates with different contrast agents was done using Nano-ZS instrument (Malvern Instruments, Worcestershire, UK) with Zetasizer software version 7.11. Samples were loaded into low volume quartz cuvette (Hellma, GmbH) and data were acquired after 300 seconds equilibration at 25°C. Dispersion Technology Software (DTS) was used for data collection and analysis. Size of conjugates was analyzed by comparing intensity, volume, and number distributions, which were used to calculate the average size.

### Enzymatic assay

PGD2 produced from the PGH2 substrate as a result of isomerase activity of functional L-PGDS was treated with MOX HCl to form a stable product. All standards and substrate solutions were prepared as per booklet instructions for Prostaglandin D2-MOX ELISA kit purchased from Cayman Chemical, USA. 1μM of purified L-PGDS or its conjugates were added to 5pg/ml, 50pg/ml, 500pg/ml, and 5000pg/ml samples of PGH2. The reaction was stopped by adding 1M HCl. Amount of PGD2 formed as a result of enzymatic response was calculated by measuring PGD2-MOX bound to plate antibody.

### Cell viability assay

Human neuroblastoma cells SH-SY5Y were tested for cell viability upon addition of L-PGDS and its conjugates using MTT (3-(4,5-dimethylthiazolyl-2)-2,5-diphenyltetrazolium bromide) colorimetric assay based on cellular metabolic activity. SH-SY5Y cells were grown in DMEM/ F-12 media with 10% FBS till 70% confluency to ensure proper morphology. MTT assay was done in 96 well plates with 2500cells per well. Cells were incubated for 24 h at 37°C after addition of samples. 10μl of MTT (5mg/ml) was added in each well and incubated for 4 h before removing excess media and dissolving the formazan crystals in 100μl DMSO for 1 h in the dark, before the measurement of absorbance at 570nm and subtracted reference at 630nm wavelength.

### Animal permit

All applicable international, national, and/or institutional guidelines for the care and use of animals were followed. All procedures performed in studies involving animals were in accordance with the ethical standards of the Institutional Animal Care and Use Committee (A*STAR Biological Resource Centre, Singapore, IACUC #161134). 1-year old Control C57Bl6 (N =4) and 3xTgAD (N = 3) mice have been used in the experiments [9]. One control and two 3xTg mice underwent intra-ventricular infusion of LA, one 3xTg mouse underwent infusion with LD conjugate. Two control mice underwent intra-nasal administration of LA, while one mouse underwent intra-nasal administration of the contrast agent AfFtnAA. Animals were kept in standard housing with 12/12 light/dark cycle with food and water provided *ad libitum*.

### Intra-ventricular infusion

Mice were anaesthetised with a mixture of ketamine/xylazine (ketamine 75 mg/kg, xylazine 10 mg/kg). The head was shaved and cleaned with three wipes of Betadine® and ethanol (70%). Lidocaine was administered subcutaneously, *in situ*. Each animal was kept on a warm pad to prevent hypothermia, and the head was positioned in a stereotaxic frame, and the protective ophthalmic gel was applied to avoid dryness. An incision was made on the skin along the anterior-posterior axis. Perforation of the skull was performed with a driller (driller tip Ø .9 mm^2^) in the predefined location. The coordinates for the craniotomy were −0.5 from bregma, +1 left from the middle line; injection was carried out at −1.5 mm from the brain surface. A 1 μl injection of LA or LD conjugate in each of the left and right ventricles was performed in the target location through a precision pump (KD Scientific Inc., Harvard Bioscience) with a 10 ul NanoFil syringe with a 33-gauge beveled needle (NF33BV-2). The needle was kept in location for 10 minutes after the injection was completed, to exclude backflow. The skin was sutured using absorbable sutures. Buprenorphine was administered post-surgically to each animal. Animal recovery took place on a warm pad.

### Intra-nasal administration

Mice were lightly anesthetized with isoflurane 1% and held vertically. 5 μl of L-PGDS conjugates or contrast agents were administered into a nostril using a pipette. After 5min, an additional 5 μl samples were administered into the second nostril, as previously reported [2].

### Magnetic resonance imaging

Mice were anesthetized with 5% isoflurane in ¼ O2 in medical air. Mice were positioned on a water-heated MR-compatible cradle. Anesthesia was delivered via a face mask using isoflurane 1.5%. Mice temperature and breathing rate were monitored during the experiments. MRI experiments on live anesthetized mice were conducted on 9.4T Bruker Biospec equipment using a 2×2 phased-array receiver coil. Paravision version 6.0 was used to run T2- and T1-weighted MSME and RARE sequences to acquire image data. Excised and fixed mouse brains were also images on 14T Bruker Avance III spectrometer with MicWB40 microimaging probe for longer imaging time to achieve higher resolution.

### Histological Staining

Mouse brains injected with LA conjugates were isolated from the animals euthanized after MRI imaging and the whole tissue was frozen in OCT medium. These brain tissues were sliced in 20μm layers using Leica cryostat. Excised slices were stores in −20°C till staining. Brain tissue was stained using Prussian blue to identify iron in nanoparticles inside the brain and congo red to stain amyloid structures. Stained slices were visualized under a light microscope. Ninhydrin reagent was used to identify conjugate location owing to the capability of detecting free amino groups. Diaminobenzidine-enhanced Prussian Blue stain was used to detect iron content in the excised brain tissue.

### Data Analysis

All the curves were plotted in Originlab 7 or GraphPad Prism. MRI and histological images were analyzed using ImageJ. MRI data was processed in Paravision 6.0.1 software.

## Abbreviations

Aβ: amyloid-β
AD: alzheimer’s disease
L-PGDS: lipocalin type prostaglandin d synthase
CSF: cerebrospinal fluid
MRI: magnetic resonance imaging
3xTg: triple transgenic
PEG: polyethylene glycol
DLS: dynamic light scattering
RARE: rapid imaging with refocused echoes
PGD2: prostaglandin d2
PGH2: prostaglandin H2
TEM: transmission electon microscopy
GRE: gradient recalled echo
AfFtnAA: ferritin mutant from from archaeoglobus fulgidus
MNS: magnetic nanostructures with peg coating
SiNP: silica coated nanoparticles
DNP: dopamine coated nanoparticles
LA: lpgds-afftnaa conjugate
LM: lpgds-mns conjugate
LS: lpgds-sinp conjugate
LD: lpgds-dnp conjugate
MTT: 3-(4,5-dimethylthiazol-2-yl)-2,5-diphenyl tetrazolium bromide
ThT: thioflavin t

## Supplementary figures and table

**Figure S1:**
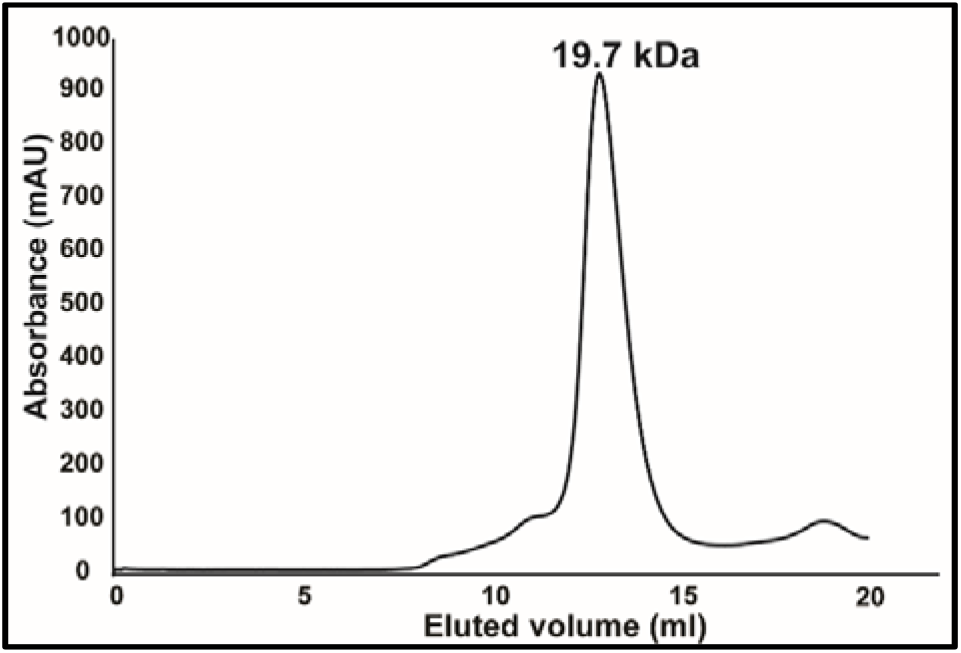
Size exclusion chromatogram for recombinantly expressed wild-type L-PGDS. Purified protein fractions collected under the peak were used for subsequent assays.

**Figure S2:**
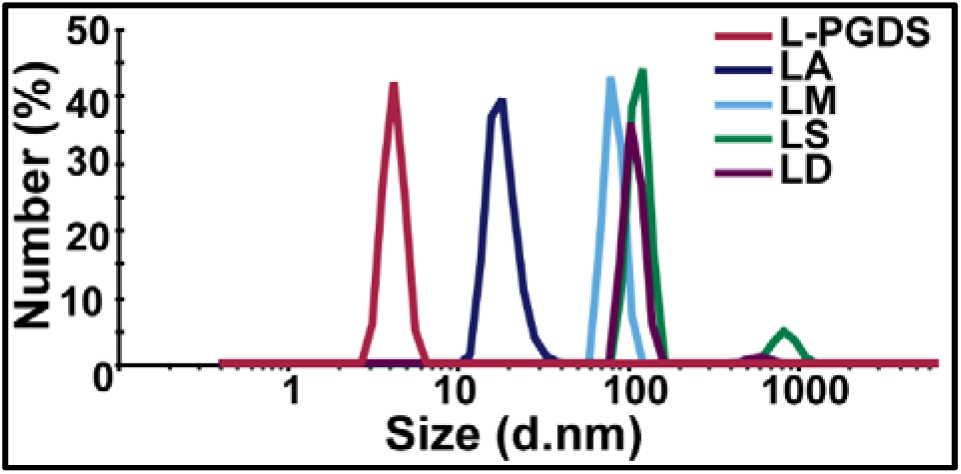
Characterization of size for the LPGDS-IONP conjugate probes. Hydrodynamic size of the conjugates was determined using Dynamic Light Scattering (DLS). Compared to the mean size diameter of L-PGDS (red) protein as 4.2 nm, conjugates LA (blue) were 20.1 nm, LM (cyan) were 68.3 nm, LS (green) were 118.6 nm, and LD (purple) were 100.8 nm.

**Figure S3:**
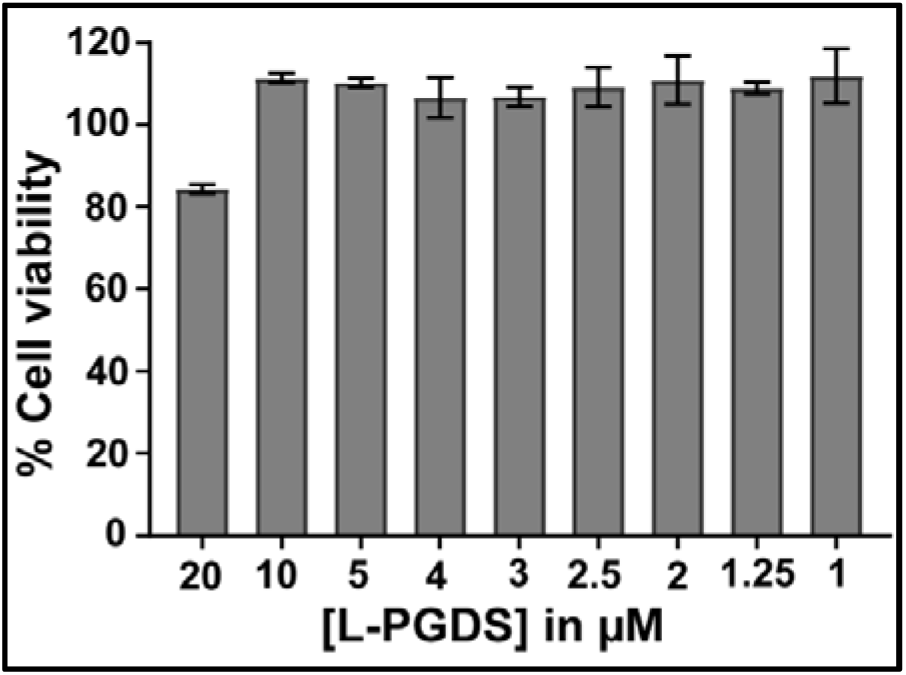
Cell viability assay of LPGDS-IONP conjugates using SHSY-5Y human neuroblastoma cells. Percentage cell viability of SHSY-5Y human neuroblastoma cells, mean ± SEM (n=3), upon incubation with different concentration of L-PGDS protein for 24 h, was calculated based on the absorbance of formazan product at 570 nm metabolized from MTT substrate.

**Figure S4:**
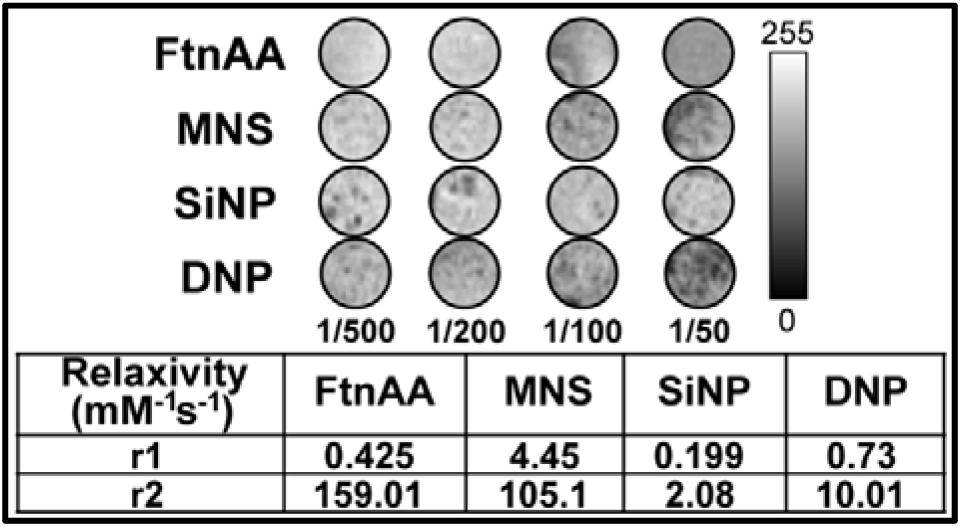
Magnetic contrast comparison for differently coated iron oxide nanoparticles (IONPs). (a) T2-weighted contrast images for IONPs. T2* images were obtained using Multiple Gradient Echo (MGE) sequence with TR: 1000 ms, TE_eff_: 55 ms, Flip angle: 50° (b) Relaxivity values for each contrast agent calculated for IONP samples using spin-echo RARE sequence with variable TR and multiple echoes, TR_max_: 15000 ms, TE: 40 ms, Rare factor: 8, FOV: 20 × 20 mm, image size: 256×256, slice thickness: 0.5 mm, no. of T1 experiments: 8, echo images: 6.

**Figure S5:**
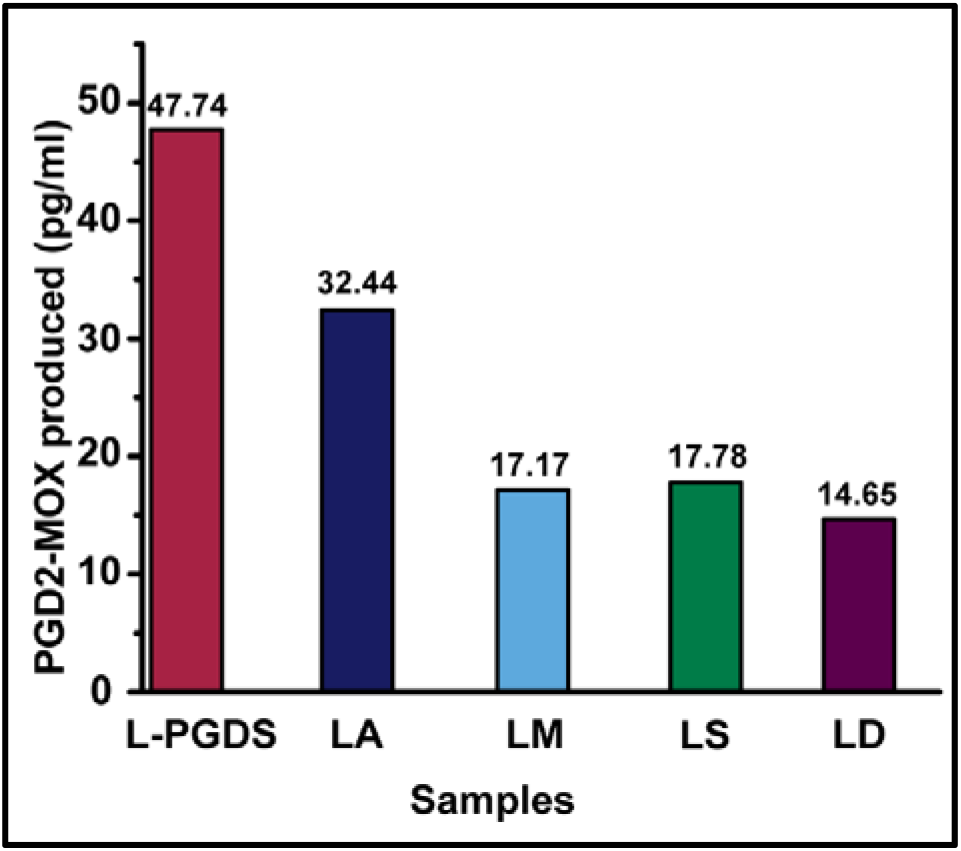
Comparison of enzymatic activity of the LPGDS-IONP conjugate probes. The enzymatic activity of native L-PGDS and the conjugates was determined using ELISA based MOX kit from Cayman Chemical. PGD2-MOX is formed as a stable derivative of the PGD2 product from L-PGDS enzymatic activity and its concentration was determined from the standard curve using four-parametric logistic fit.

**Figure S6:**
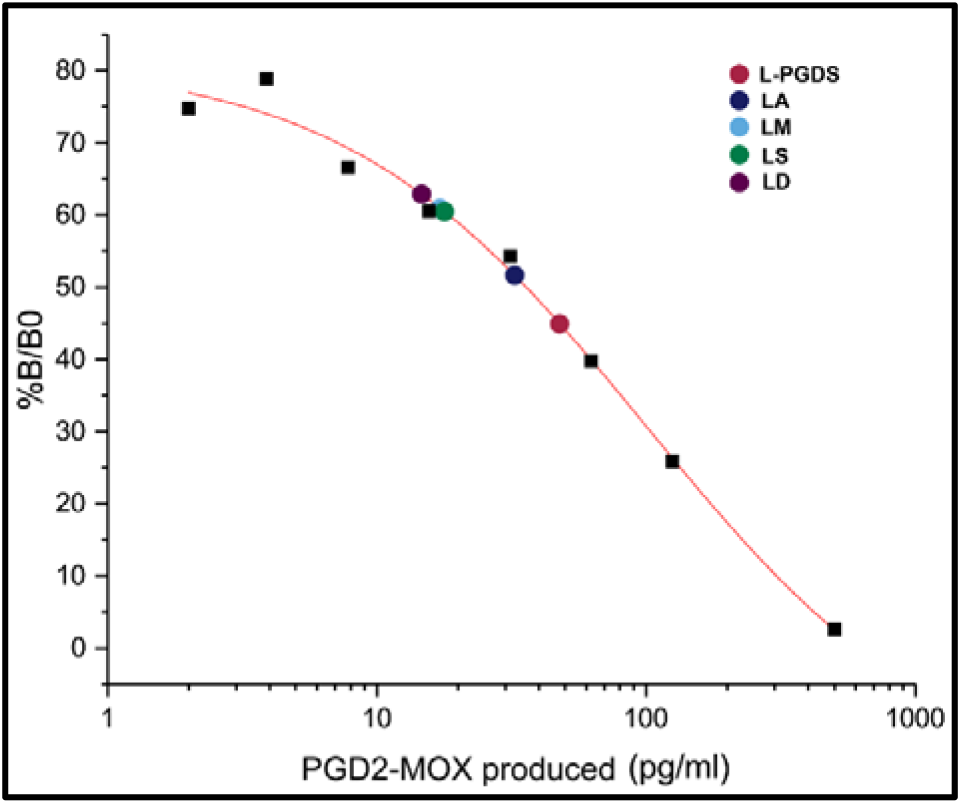
A 4-parameter logistic curve fitted for standard values based on the standard sample absorbance readings at 410nm as per manufacturer’s instructions. The standard curve was used for the concentration determination of the PGD2-MOX product.

**Figure S7:**
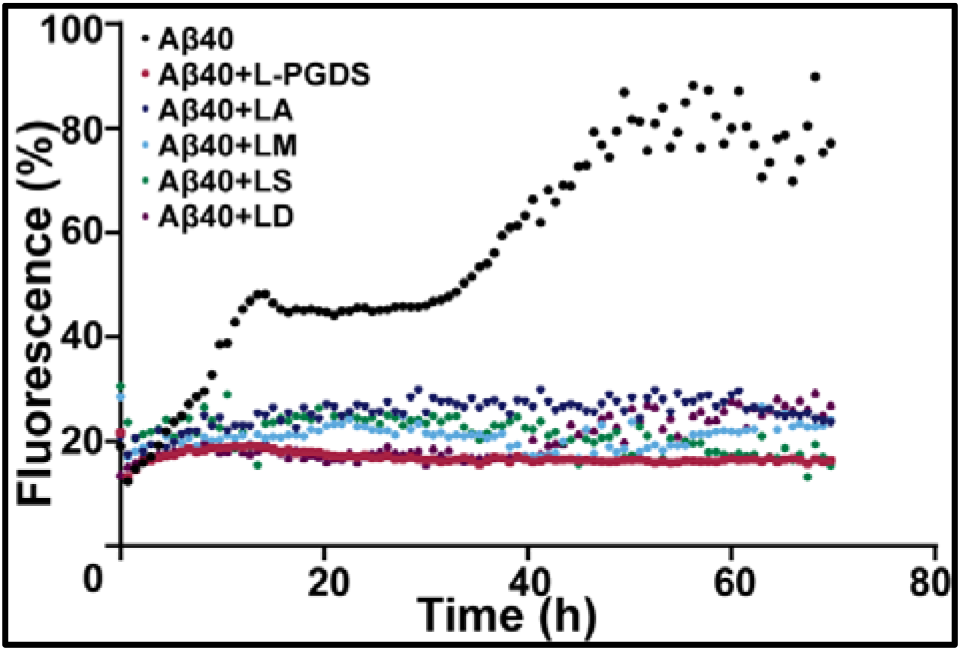
Inhibition of Aβ aggregation by LPGDS-IONP conjugates. Thioflavin T fluorophore intensity (Ex-445 nm, Em-485 nm) of 50 μM Aβ (1-40) monomers in the absence (black) and presence of 2.5 μM protein as native L-PGDS (red) and in conjugated probes (LA-blue, LM-cyan, LS-green, and LD-purple) was measured every 15 min with shaking at 37°C for 72 h.

**Figure S8:**
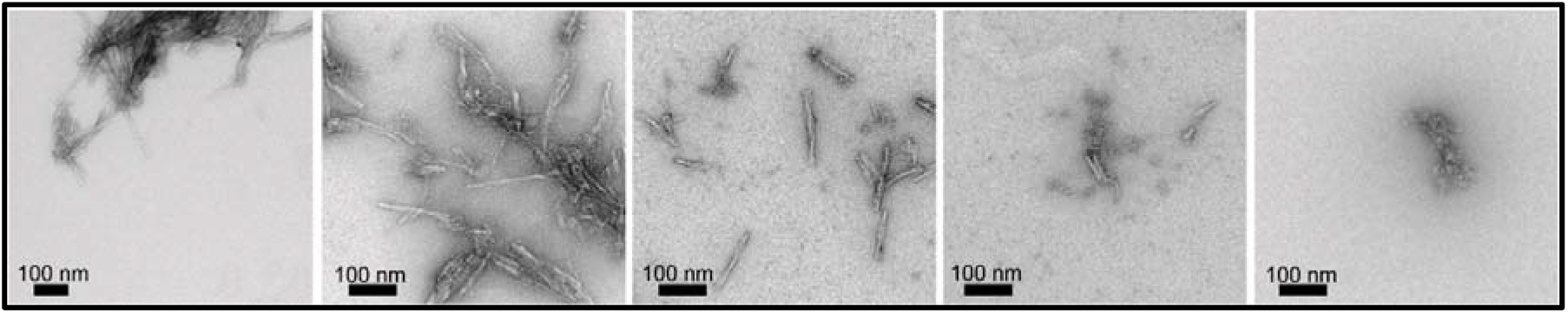
Transmission electron microscopy of Aβ fibrils incubated with LA conjugates for 30 min to visualize breakdown of the Aβ (1-40) fibrils.

**Figure S9:**
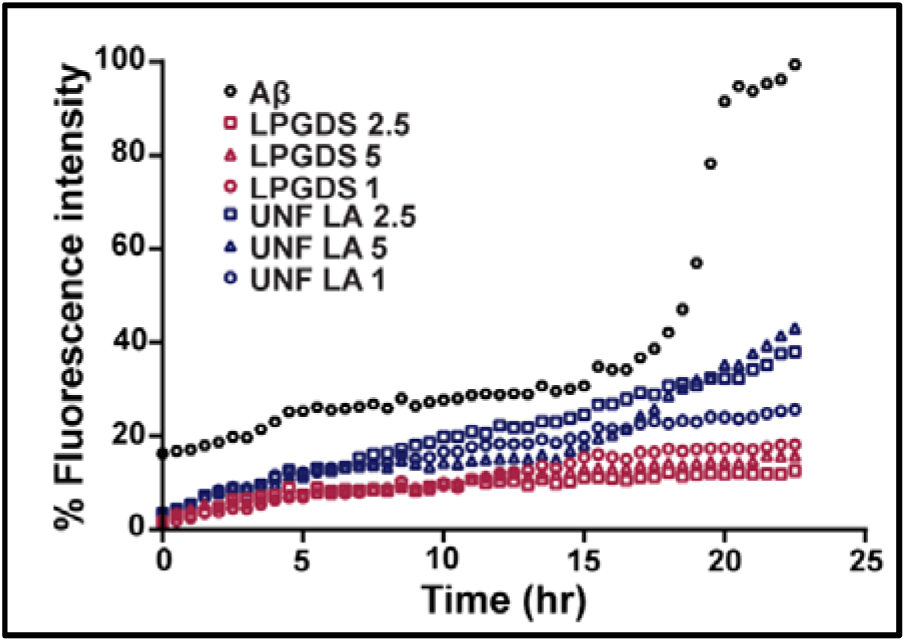
Characterization of inhibition of amyloid aggregation in presence of L-PGDs conjugates. Thioflavin T fluorophore intensity (Ex-445 nm, Em-485 nm) of Aβ (1-40) monomers in the presence of equal concentrations of L-PGDS and LA conjugates was measured for 72 hrs as a correlation to their inhibition of aggregation.

**Figure S10:**
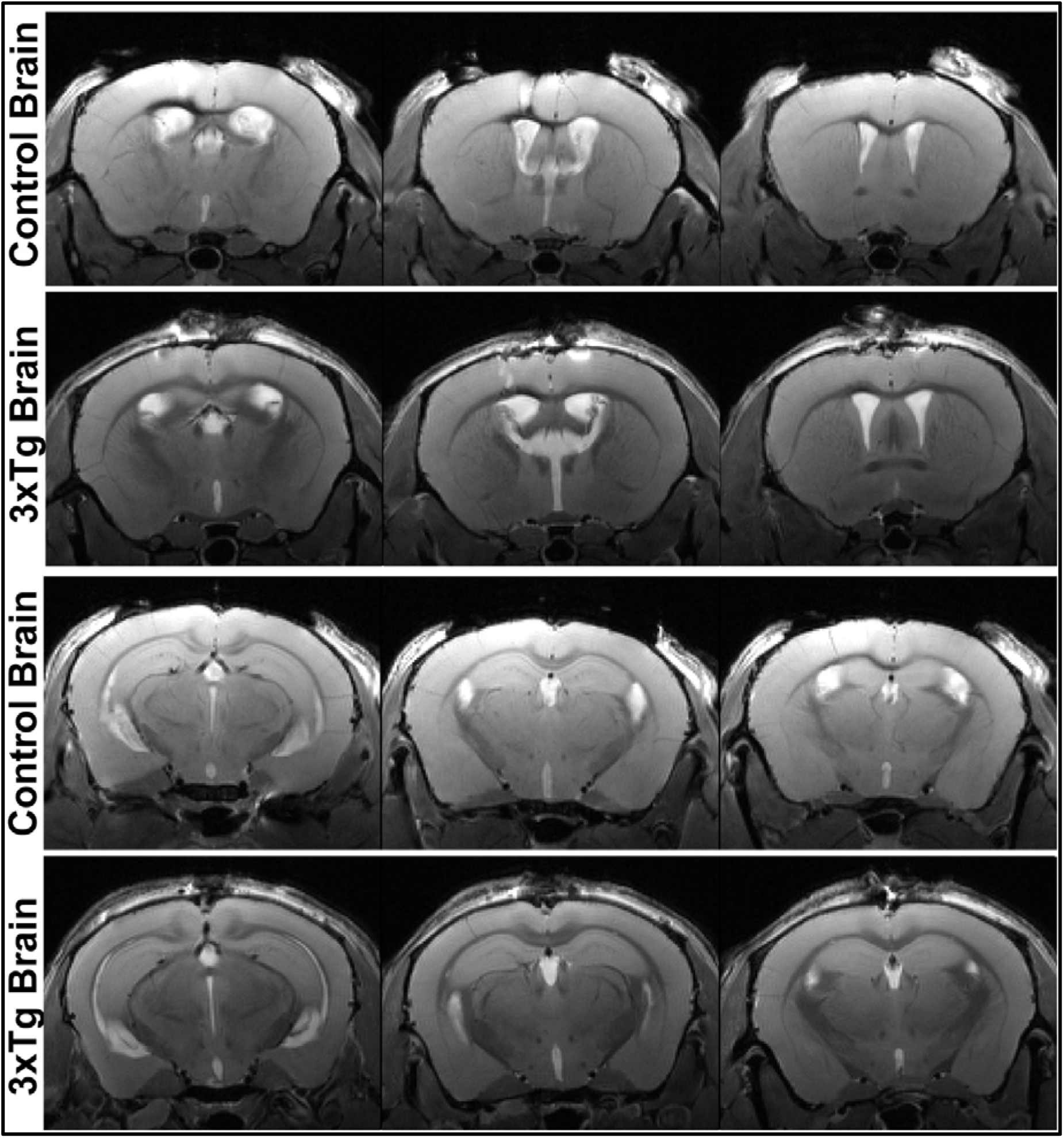
Contrast comparison upon intraventricular injections of LA conjugates in healthy versus AD mice brain. Enhancement of T2* contrast in 1-year old 3xTg mouse brain compared to age-matched control mouse brain. The increase in hypointensity is marked by red arrows in the 3xTg brain. The coronal images were acquired at 11.4 T using T2_TurboRARE sequence with image size was 200 × 106 pixels, with field of view 17 × 9 mm, RARE factor: 4, TE 15.56 ms, TR 3500 ms with38 slices of thickness 0.4 mm each through the mouse brain, scan time: 3 min 2 sec, motion averaging, flip back, fat suppression.

**Figure S11:**
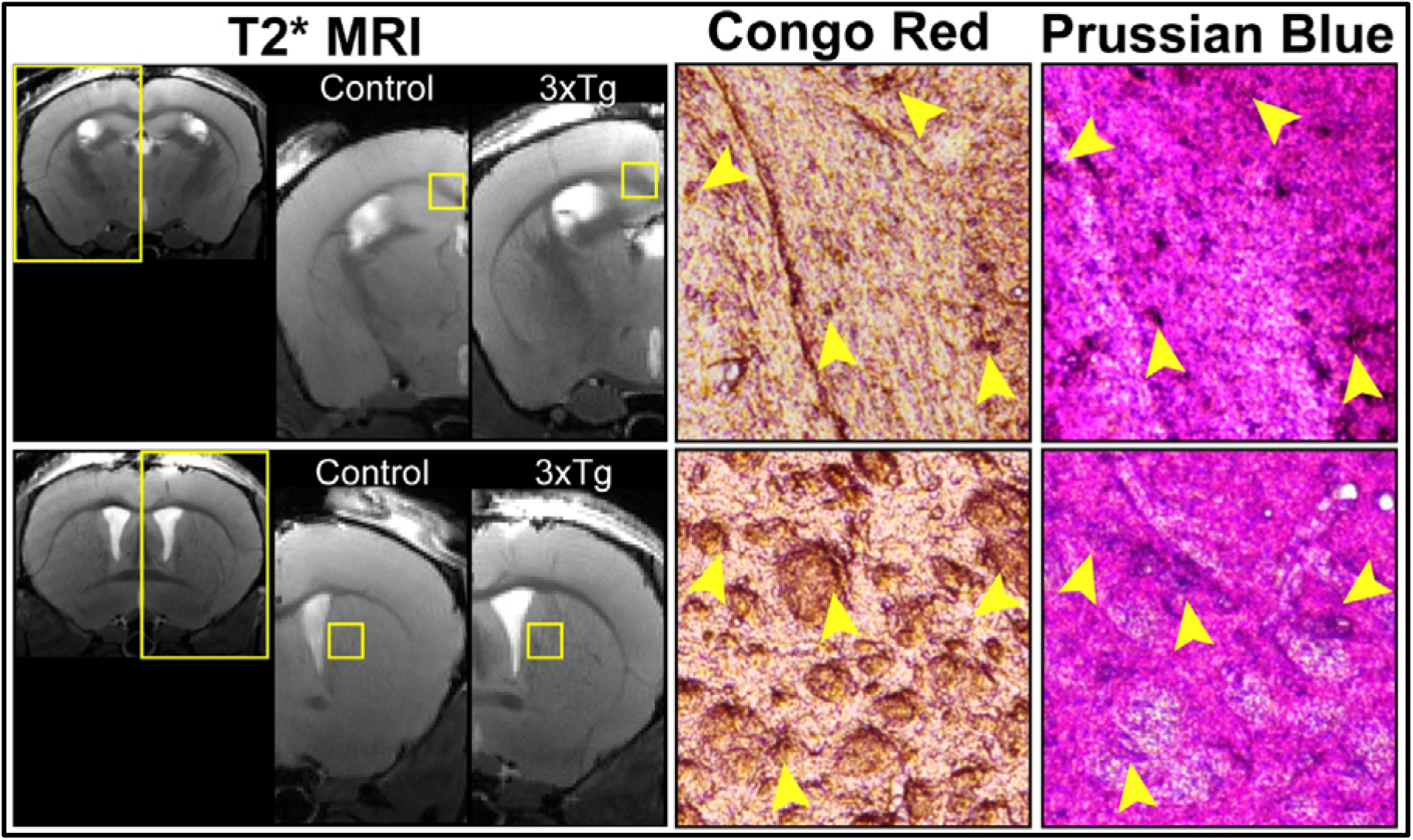
Histological staining of 3xTg AD brain slices post intraventricle injections. Regions of hypointensity in T2* MRI images are stained with Congo red and Prussian blue staining of mouse brain slices of different regions after excision from 3xTg mice 72h post intraventricular injection with LA conjugates.

**Figure S12:**
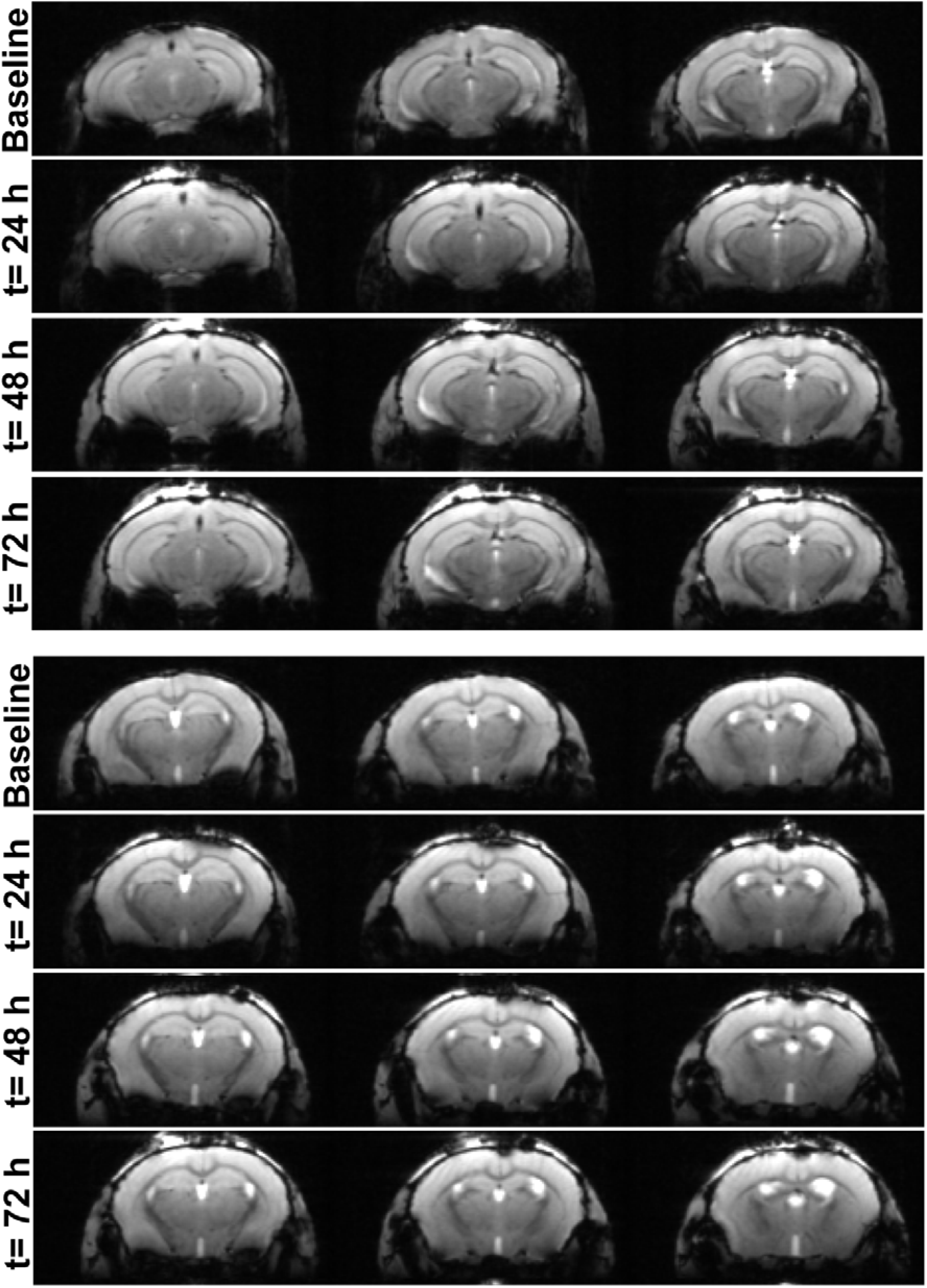
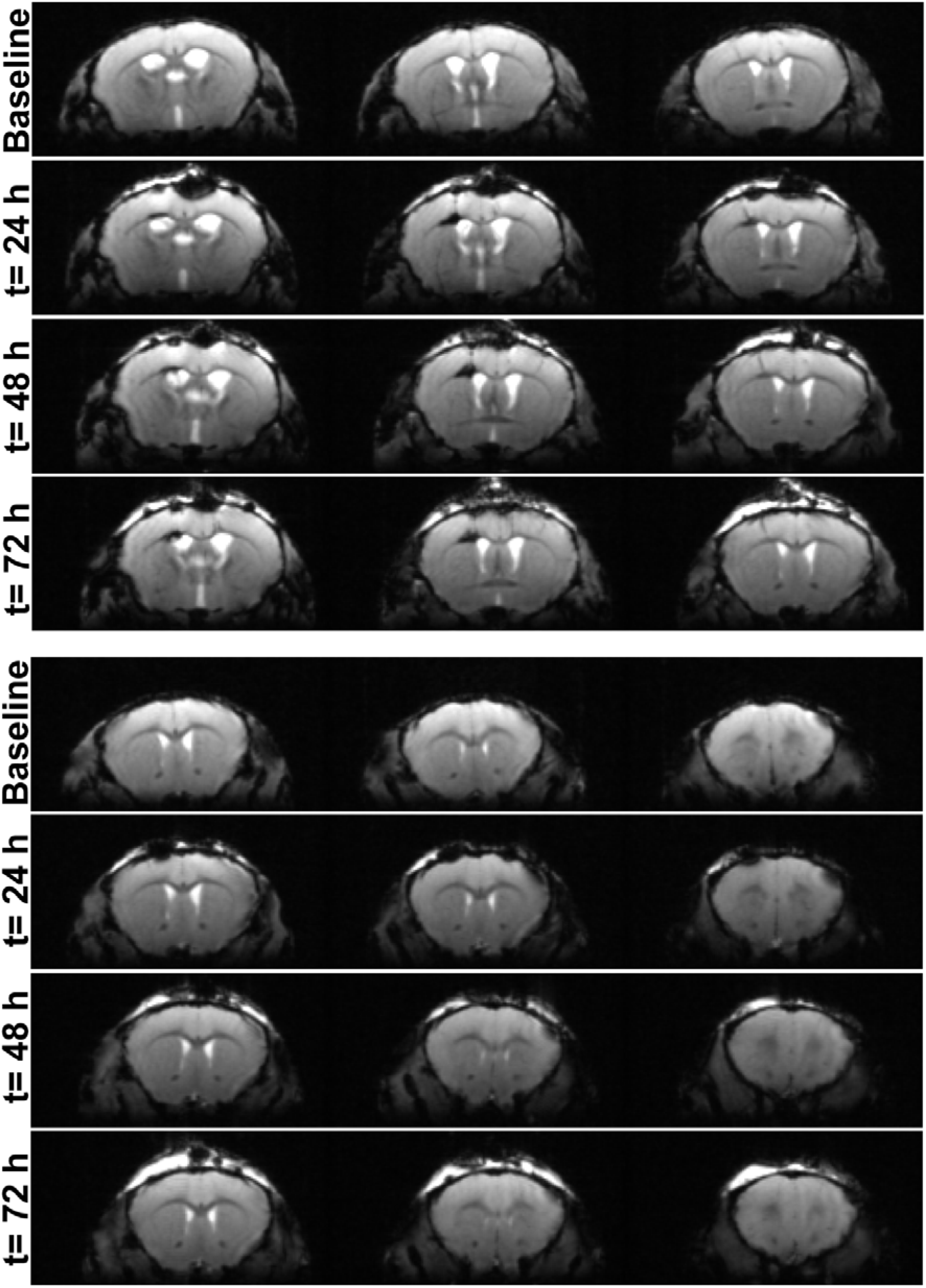
Comparison of T2* contrast enhancement in the 3xTg AD mouse post intraventricular injection of LA conjugates. Regions of hypointensity and susceptibility variation due to the iron deposits in the mid-brain slices seen in the images 24 h, 48 h, and 72 h post intraventricular injection images have been marked with red arrows. The coronal images were acquired at 11.4 T using T2*-FLASH sequence with an image size 200 × 106, field of view 17 × 9 mm, Flip angle 60°,TR: 1500 ms, TE: 10 ms, for 38 slices with thickness 0.4 mm each through the mouse brain; scan time: 6 min, motion averaging, fat suppression.

**Figure S13:**
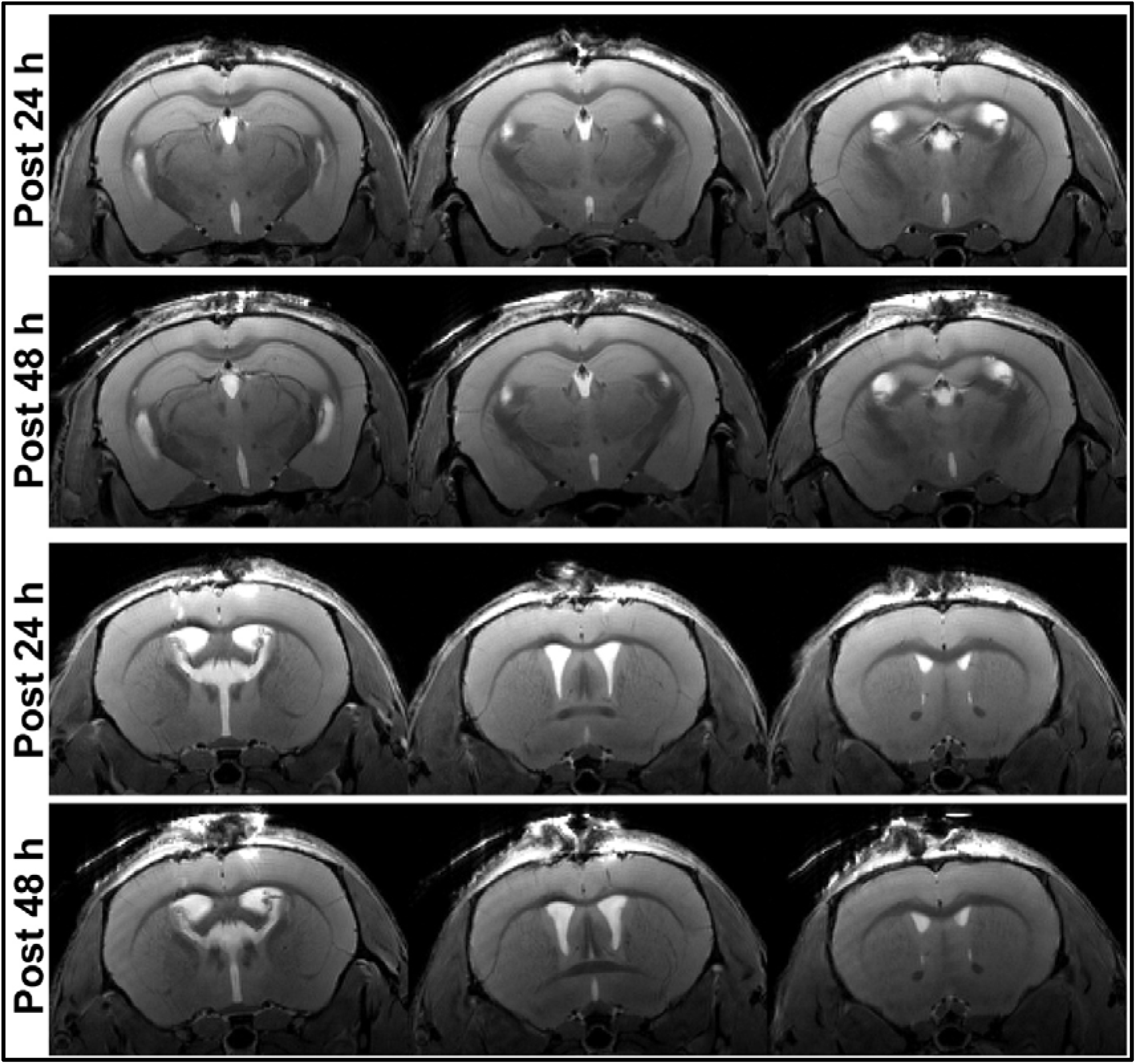
MRI of AD mouse brain post 24 and 48 h of intraventricular injection of LA conjugates in 3xTg mouse brain. The contract/regions of accumulations of the particles upon diffusion is shown with red arrows.

**Figure S14:**
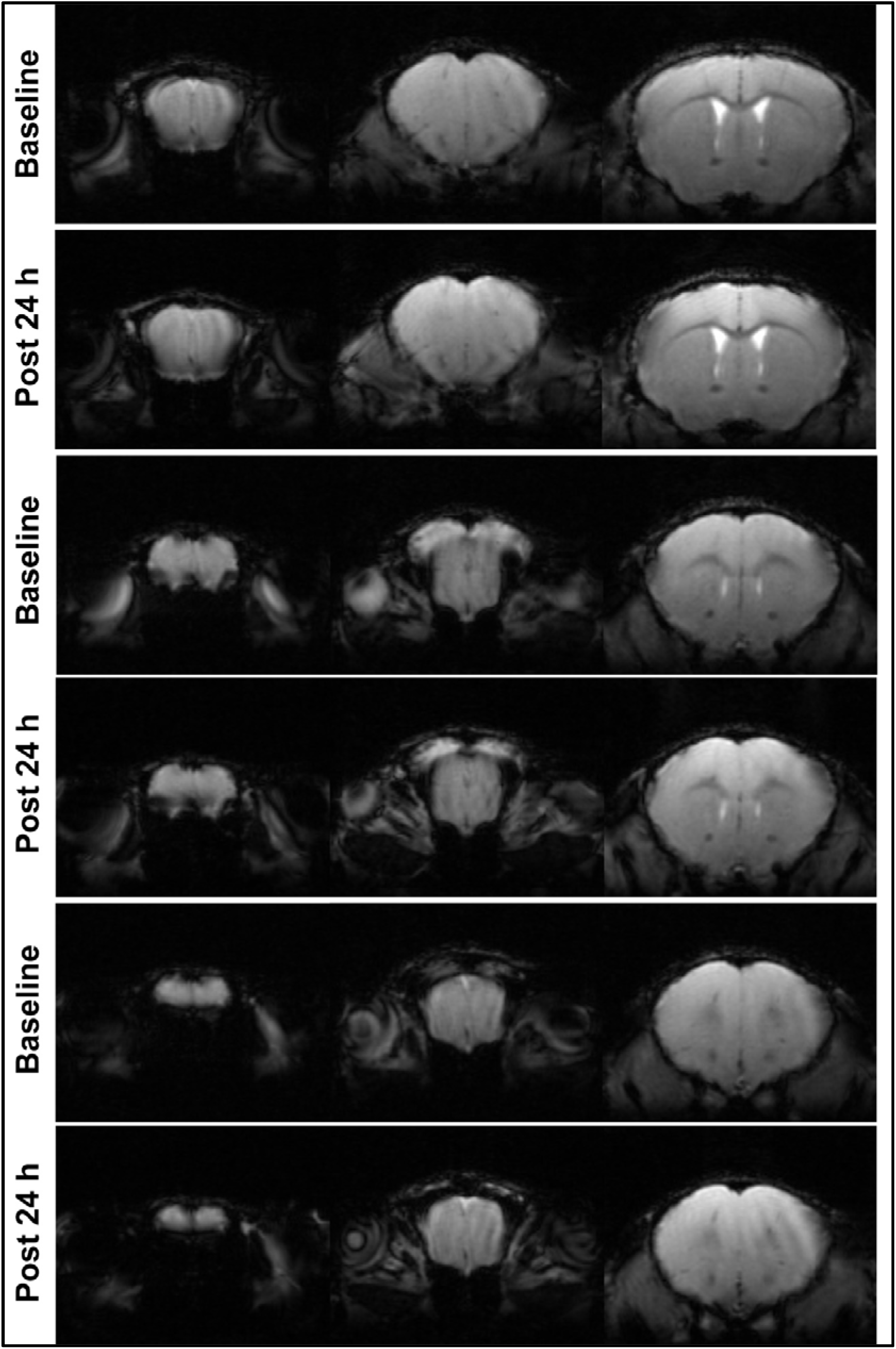
Brain detection of conjugates in control mice by intranasal administration. The left and right panels represent the same slice in T2*-weighted FLASH images taken before and 12 hours after the intranasal delivery of LA conjugates in the healthy mice. The coronal images were acquired at 11.4 T using MSME scans with image size was 200 × 106 pixels, with field of view 17 × 9 mm, 60° flip angle, TE 10ms, TR 1500 ms with 3 averages each for 38 slices through the mouse brain.

**Figure S15:**
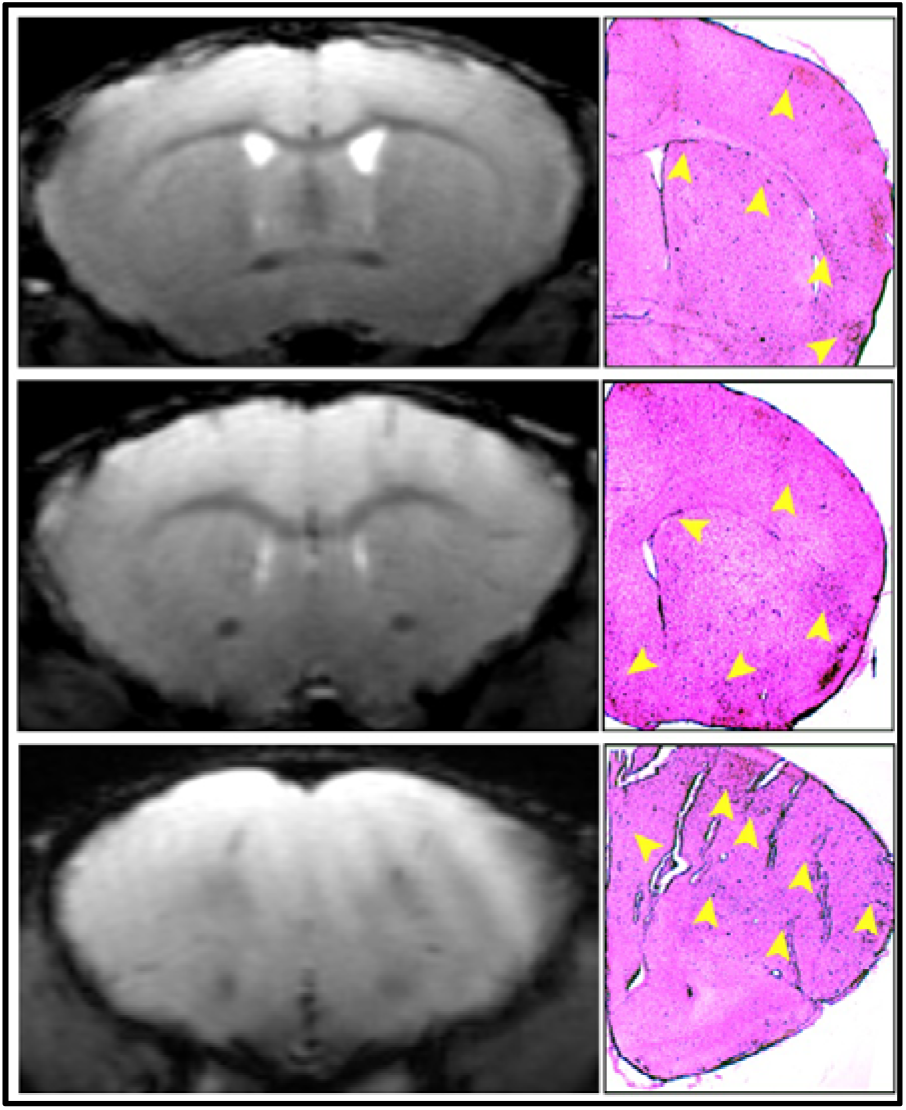
Detection of LA conjugates in healthy mice brain post intranasal administration. The conjugates in the brain were detected using T2* contrast MRI and Prussian blue staining of the brain slices. The presence of LA conjugates leads to hypointensity in MRI images and blue coloration in the stained slices.

**Table S1:** T2* values (ms) for select brain regions of interest before and 72 h after intraventricular injection of LA conjugates in 3xTg AD mouse. Numbers were calculated from selected circular regions of interest in T2 map_MSME images with TR: 2200 ms, echo spacing: 8.16 ms, image 80 × 80, FOV: 20 × 20 mm, slice thickness: 0.5 mm, no. of echo images: 9, flip back, excitation 90 degree, refocusing 180 degree.

| **0 h** | **72 h** | **Brain Region** |
| --- | --- | --- |
| 36.14 ± 0.45 | 36.77 ± 0.57 | Olfactory bulb |
| 37.21 ± 0.63 | 36.69 ± 0.60 | Neocortex |
| 36.14 ± 0.80 | 35.08 ± 0.52 | Corpus callosum |
| 36.02 ± 0.50 | 35.70 ± 0.53 | Corpus callosum |
| 38.03 ± 1.09 | 36.01 ± 0.61 | Hypothalamus |
| 36.54 ± 1.02 | 33.41 ± 0.50 | Thalamus |
| 35.42 ± 1.16 | 34.63 ± 0.77 | Hippocampus |
| 33.24 ± 0.70 | 32.85 ± 0.68 | Cerebral aqueduct |
| 34.45 ± 0.68 | 33.73 ± 0.72 | Cerebellum |

